# Modelling the effectiveness of Integrated Pest Management strategies for the control of Septoria tritici blotch

**DOI:** 10.1101/2025.02.28.639837

**Authors:** Elliot M.R. Vincent, Edward M. Hill, Stephen Parnell

## Abstract

Reducing reliance on pesticides is an important global challenge. With increasing constraints on their use, in recent years there has been a declining trend in pesticide use for arable crops in the UK. But with increasing disease pressures and global demand for food, there is a greater need for effective measures of pest and disease control.

These circumstances highlight the need for widespread adoption of sustainable alternative control measures. Integrated Pest Management (IPM) is one such solution, comprising a set of management strategies which focus on the long-term prevention, detection and control of pests, weeds and diseases. While many of these methods are acknowledged to offer effective control, their implementation has thus far been limited in practice.

As a case study we considered Septoria tritici blotch (STB) (*Zymoseptoria tritici*), an economically important disease of wheat. We used epidemiological modelling techniques to investigate the potential of different IPM control strategies (crop residue burial, delayed sowing, variety mixtures and biocontrols). Combining existing data with a deterministic, compartmental infectious disease model of STB transmission, we simulated the implementation of an IPM regime into the STB disease system. We investigated the outcomes on disease prevalence and crop yield when comparing conventional and IPM control regimes. In a single field, for the individual implementation of IPM measures we found the starkest difference in potential yield outcomes between delayed sowing and biocontrols (greatest yields), and crop residue burial and variety mixtures (lowest yields). We also found that the collective use of IPM measures has the potential to offer individual growers comparable control to a standard fungicide regime. For a multi-field setting, representing a community of crop growers, a high proportion of growers using IPM can reduce the level of external infection incurred by the growers who maintain a fungicide regime.

**Author Summary:** With the UK Government seeking to reduce the environmental risk posed by pesticides, the agricultural industry is under increasing pressure to explore alternative methods of disease control. One such alternative method is Integrated Pest Management (IPM), which consists of a variety of management strategies for long-term prevention, detection and control of crop diseases. In our study, we simulate the potential outcomes of using IPM for the control of Septoria tritici blotch, a common disease of wheat. Our results suggest that a regime of IPM control methods may offer growers comparable yields and disease control to conventional fungicide treatments. Furthermore, in a wider system of crop growers, a higher proportion using IPM can reduce the level of infection incurred by all growers in the system, including those who do not use IPM. These findings can offer insight to crop growers who are considering the use of IPM, and to policy-makers who are interested in encouraging its uptake, by validating and quantifying its effectiveness relative to current standard practices.

## 1 Introduction

Global demand for food is widely projected to increase in the coming decades, with many predicting a rise between 35% to 56% from 2010 and 2050 [1]. The agricultural industry faces the need to increase its production, while also working against mounting pressure from plant pests and diseases, which are currently estimated to be the cause of up to 40% yield losses globally [2]. One such disease is Septoria tritici blotch (STB) (commonly called ‘septoria’, ‘septoria leaf blotch’), a foliar disease that primarily affects wheat, whose causative agent is *Zymoseptoria tritici*. The UK’s Agriculture and Horticulture Development Board (AHDB) report STB as the most important and damaging foliar disease on winter wheat [3]. The pathogen causes necrotic lesions to develop on the leaves, reducing the area of healthy green tissue available for photosynthesis, resulting in yield loss and reduced grain quality. AHDB state that crops severely affected by STB can suffer losses of up to 50% [3].

To combat these threats to crop health, chemical control remains a staple of conventional crop protection. However, due to the negative ecological and environmental consequences of their use [4], there have been increasing constraints on pesticide use for pest and disease management (e.g. removal of some chemical pesticides from UK markets). These constraints have resulted in a recent declining trend in pesticide use on arable crops in the UK. The ‘Arable crops in the United Kingdom’ survey for 2022 reported, compared to 2018, a decrease in area treated with pesticides of 6% and a decrease in weight of pesticide applied of 14% [5]. These circumstances bring to the fore the need for widespread adoption of sustainable alternative solutions for crop pest and disease management.

Much focus for a sustainable alternative has been put on Integrated Pest Management (IPM). IPM is not a single control method, but a programme of coordinated, environmentally-sensitive control measures that can be used to prevent, detect and control crop pests or diseases [6]. It can comprise a wide range of potential strategies, such as the use of resistant varieties, biological controls, modification of sowing practices, and the targetted and controlled use of chemical pesticides. One of the main, generally agreed-upon, aims of IPM is to reduce reliance on the use of synthetic pesticides [7]. Nevertheless, despite long-term empirical evidence of the effectiveness of IPM, its adoption in practice has been limited. Growers face barriers to uptake in the form of access to existing tools and knowledge, risk, ease of implementation, and cost [8].

There remain several unanswered questions regarding IPM and its use in the control of STB. Previous modelling work on STB has investigated the rate of development of fungicide resistance under the use of high- or low-risk fungicides [9], the use of fungicide mixtures and alternation [10–12], and the timing of fungicide applications [13]. One study, by Taylor and Cunniffe [14], explored the use of resistant cultivars in combination with fungicide treatments, which would be considered an IPM strategy commonly in use by growers in the UK. Previous modelling work has not, however, investigated the use of IPM controls that are presently available to farmers but not commonly deployed in practice. Examples of such IPM controls that have had low uptake to date in the UK are variety mixtures, residue management, varied sowing dates, and biocontrols [15].

A crucial benefit that modelling can offer is to complement other types of empirical study. It would be ideal to test the effectiveness of IPM regimes and compare them to current standard fungicide practices by performing real-world, long-term trials. Such studies, however, can be challenging to perform due to operational and economic barriers. Alternatively, modelling can provide a means to gain initial insight into such questions, as has been done previously with a number of different diseases and management strategies [16–18]. Nevertheless, at the time of writing we have not identified studies that have used modelling to contrast the possible impacts on yields of an IPM control strategy versus a conventional fungicide strategy. Moreover, such work can help identify where additional data to reduce parameter uncertainty can improve the robustness of model predictions. In this paper, we use modelling to address the following questions (with respect to yields):

- How does an individual IPM measure perform compared to either having no control measures, or using conventional fungicide treatment?
- Can the collective use of multiple IPM measures outperform a conventional fungicide regime?
- Considering a community of crop growers, is there an optimal balance of IPM farms with conventional fungicide farms?

To provide insight into these questions, we present an infection transmission model for STB whose novel contribution is its incorporation of IPM interventions. We initially acquired from the available literature plausible value ranges for the model parameters. We then performed model simulations to investigate the potential impact on yield of chemical control (i.e. fungicide use) and four different IPM control strategies (crop residue burial, delayed sowing, variety mixtures and biocontrols). From a single-field perspective, we found that collective use of multiple IPM measures has the potential to offer individual crop growers comparable control to a standard fungicide regime. For a multifield system, we show circumstances where the presence of IPM-using fields can benefit all growers by boosting yields. By highlighting the potential for a ‘free rider’ situation (those using a chemical control regime benefitting from others using IPM), we anticipate our findings to be a starting point towards a combined epidemiological-behavioural model for growers’ intentions towards disease control. The framework we present is also adaptable to other plant pathogens, allowing researchers to investigate the implications of IPM usage on infection burden and yield outcomes for other crops.

## 2 Methods

To carry out our investigation, we first developed a mathematical model of STB infection dynamics that included fungicide and different IPM strategies as possible controls. We summarise here the core methodological components of our study. We begin with a description of our base STB model that had no controls (Section 2.1), including how we sourced plausible model parameter values from the literature, followed by how we computed yield from a single growing season (Section 2.2). We then explain how we incorporated fungicide control into the model and its parameterisation (Section 2.3). Similar details are provided for different IPM control strategies (Section 2.4). Finally, we outline the model simulations that we performed to explore our stated research questions (Section 2.5).

### 2.1 Base STB transmission model (with no controls)

We base the transmission dynamics of STB on the model framework presented by Hobbelen *et al*. [9] (Section S1), while using the up-to-date parameter values found by Corkley *et al*. [19] (Table 1). This model is a system of ordinary differential equations (ODEs) describing the host-pathogen system of *Triticum aestivum* (winter wheat) and *Zymoseptoria tritici* (the causal organism of STB). The pathogen is not explicitly featured in the model, which instead describes the relative area of leaf tissue in each disease state (Fig. 1). Note that the model by Hobbelen *et al*. [9] deals with resistance to fungicide, which means that it features two strains of infectious spores that are resistant and susceptible to fungicide, respectively. Our model assumes the presence of only the strain that is susceptible to fungicide. The Hobbelen *et al*. [9] model uses degree-day time, fitting a sinusoidal function for degree-days per calendar day to average temperature data recorded between 1984 and 2003 in Cambridgeshire, UK. In adapting this model to the parameters and growth stage timings found by Corkley *et al*. [19], we refitted the sinusoidal temperature profile function so that the relevant wheat growth stage timings aligned with the conventional calendar dates given by HGCA [20]. We define *T*_[*X*]_ as the degree-day time when the plant reached growth stage [X] (‘GS[X]’). Throughout our analysis we considered the system behaviour over one growing season.

**Table 1.** STB transmission model and fungicide parameter values. For these parameters we used the same values as used by Corkley *et al*. [19]. We use the notation ‘dd’ to represent degree days.

| Parameter | Definition | Value |
| --- | --- | --- |
| **Plant growth stages** |  |  |
| $T_{\text{emerge}}$ | Time of GS31 (emergence of leaf 3) | 1396 dd |
| $T_{32}$ | Time of GS32 | 1495 dd |
| $T_{39}$ | Time of GS39 | 1653 dd |
| $T_{61}$ | Time of GS61 (senescence/harvesting begins) | 1891 dd |
| $T_{87}$ | Time of GS87 (senescence/harvesting ends) | 2567 dd |
| **Plant growth parameters** |  |  |
| $G$ | Host growth rate | $8.2 \times 10^{-3} \text{ dd}^{-1}$ |
| $k$ | Host carrying capacity | 4.438 |
| **Disease system parameters** |  |  |
| $\beta$ | Infection rate | $2.11 \times 10^{-2} \text{ dd}^{-1}$ |
| $\gamma$ | Rate of becoming infectious | $1/350 \text{ dd}^{-1}$ |
| $\mu$ | Rate of recovering from infection | $1/600 \text{ dd}^{-1}$ |
| $\nu$ | Decay rate of primary inoculum | $8.97 \times 10^{-3} \text{ dd}^{-1}$ |
| $\Psi$ | Initial infected leaf area in lower canopy ($P(0)$) | $1.44 \times 10^{-2}$ |
| **Fungicide parameters** |  |  |
| $\omega$ | Fungicide maximum effect | 1 |
| $\theta$ | Fungicide dose response curvature | 7.68 |
| $\delta$ | Fungicide decay rate | $1.86 \times 10^{-2} \text{ dd}^{-1}$ |

**Fig. 1.**
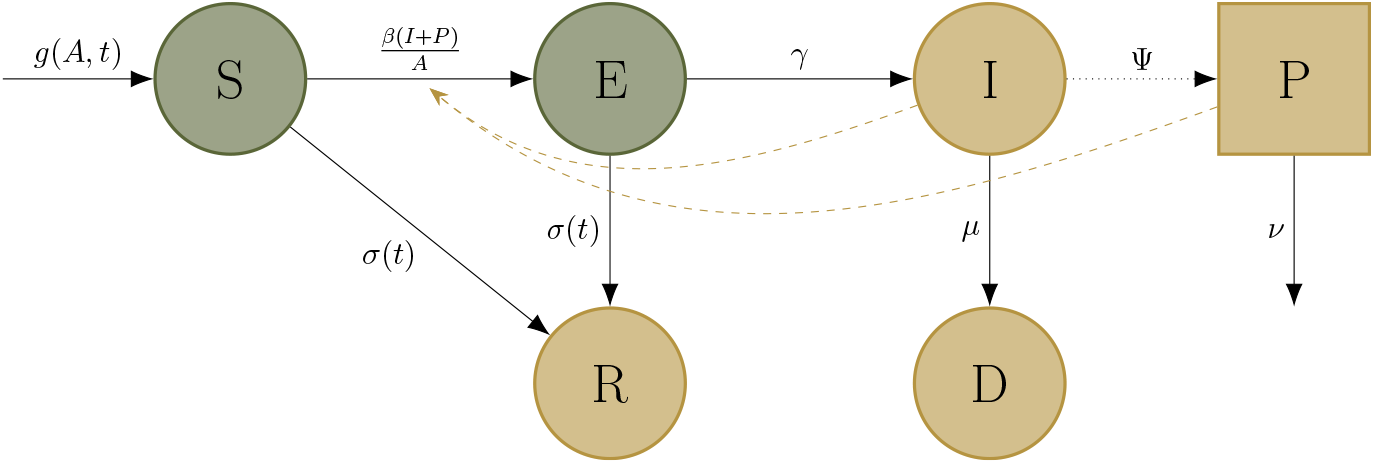
Schematic for the base STB dynamics model presented by Hobbelen *et al*. [9]. Circular nodes represent the states leaf area in the upper canopy can occupy. The square node (*P*) represents infectious leaf area in the lower canopy. Solid black lines represent the progression between states, labelled with the corresponding rates. The dotted black arrow shows between-season dynamics: infectious lesions deposit inoculum which initiates infection in the lower canopy the next year. Green nodes represent the states containing green leaf area: a healthy, uninfected state (*S*), and a latently infected state (*E*). Leaf area in these states are capable of photosynthesis, and contribute towards the yield computation. Later in the season these states experience natural senescence (*R*). Yellow nodes represent states that do not contribute to the yield computation. Infectious leaf area in the upper canopy (*I*) eventually dies due to infection (*D*), and infectious leaf area in both the upper and the lower canopy (*I* and *P*) transmit infection to healthy tissue, represented by yellow dashed arrows.

Our model has six disease states, each measured in *m*^2^ of leaf area per *m*^2^ of ground (the leaf area index). Although this is by definition a density, for simplicity it is referred to hereafter as the ‘leaf area’ because the ground area is fixed. Five of the disease states are related to the upper canopy leaves. Specifically, they describe the area of susceptible leaf tissue (labelled *S*), area occupied by latent (*E*) and infectious (*I*) disease lesions, the total leaf area that has died due to infection (*D*) and the total leaf area that has naturally senesced (*R*). Only tissue in the *I* class can transmit infection and only tissue in the *S* class can receive infection. We define the total leaf area, denoted by *A*, as *S* + *E* + *I* + *R* + *D*.

The sixth (and last) disease state, labelled *P*, represents the area of infectious lesions on the lower canopy of the plant. As disease in the upper canopy is typically initiated by infection with spores from the lower canopy, we include this lower canopy infection compartment to provide the initial infection to the plant. Infection in the lower canopy is initiated by inoculum from the previous year [11]. As defined by Hobbelen *et al*. [9] (with parameter values again taken from Corkley *et al*. [19]), we assumed infection in the lower canopy has an initial condition of Ψ, and a fixed rate of exponential decay *ν*.

The model includes time-dependent transition rates for natural growth (*g*(*A, t*)) and senescence (*σ*(*t*)) of healthy leaf tissue, and constant rates of progression from latency to infectiousness (*γ*) and death from infection (*µ*). The force of infection term is comprised of transmission from both the *I* and *P* classes, in the form of 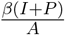.

### 2.2 Yield

We considered the yield to be proportional to the healthy area duration (HAD) over the period of grain filling; defined to be between GS61 and GS87 [9]. Given our assumption of latently infected tissue still being able to photosynthesise, we assumed the annual relative yield in our system to be

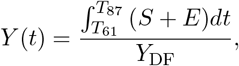

where *Y*_DF_ is the yield obtained in the disease-free scenario (i.e. a fully healthy crop) [9, 11].

### 2.3 Summary of fungicide control

In our main analyses, we assumed that growers who used fungicide did not use any additional control methods. Although this is a simplifying assumption, it allows us to more distinctly contrast the two types of control regime (fungicide versus IPM). The existing STB disease dynamics model presented by Hobbelen *et al*. [9] also included control with fungicide. To incorporate fungicide control into our model, we used functions of an analogous form to [9], but re-fit the product-dependant fungicide parameters to newer data on fungicide performance in winter wheat. Specifically, we fit to the dose response curve from 2022-2024 data for the product Revystar XE [21]; an SDHI and azole mixture that represents current standard practices for STB of wheat in the UK. In alignment with both the fungicide trial design [21] and the Hobbelen *et al*. [9] model, we assumed that fungicide treatment was applied at GS32 and GS39. We modelled fungicide concentration over time by

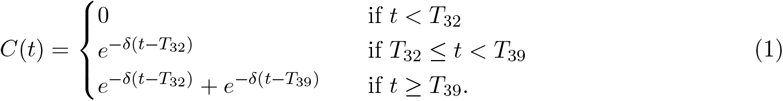

The parameters *ω, θ* and *δ* represent the maximum effect of fungicide, fungicide dose response curvature, and fungicide decay rate respectively. The values for these parameters are listed in Table 1. As *θ* and *δ* are product-dependant, these parameters were refitted, while *ω* = 1 was chosen to reflect the default dose used being the full label dose. The effect of this fungicide on both transmission, and on infection period, depends upon its concentration in the form of

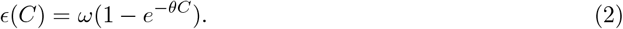

Therefore, the system of equations that govern STB infection in a field under a fungicide regime (denoted with the subscript *F*) is

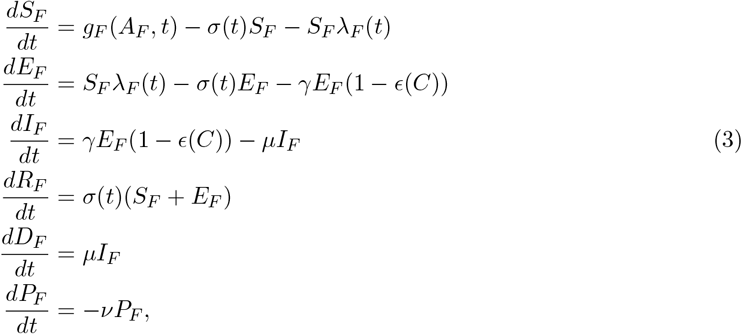

where 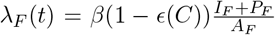, and *g*_*F*_ is identical to the function *g* in the base system [9] (Section S1). Parameter values, *σ*(*t*), and initial conditions are the same as in the base system (Table 1 and Section S1).

### 2.4 Summary of considered IPM controls

We considered the IPM measures described in the 2021 AHDB Research Review on *Enabling the uptake of integrated pest management (IPM) in UK arable rotations*, which identifies nine IPM measures applicable for the prevention of STB in wheat [15]. Based on the existing research literature, the review assigned scores on a scale of 1-5 to each measure, across a number of qualities (Fig. 2). These qualities include the effectiveness of control, strength of evidence, economic factors, implementation factors, and current and potential use.

**Fig. 2.**
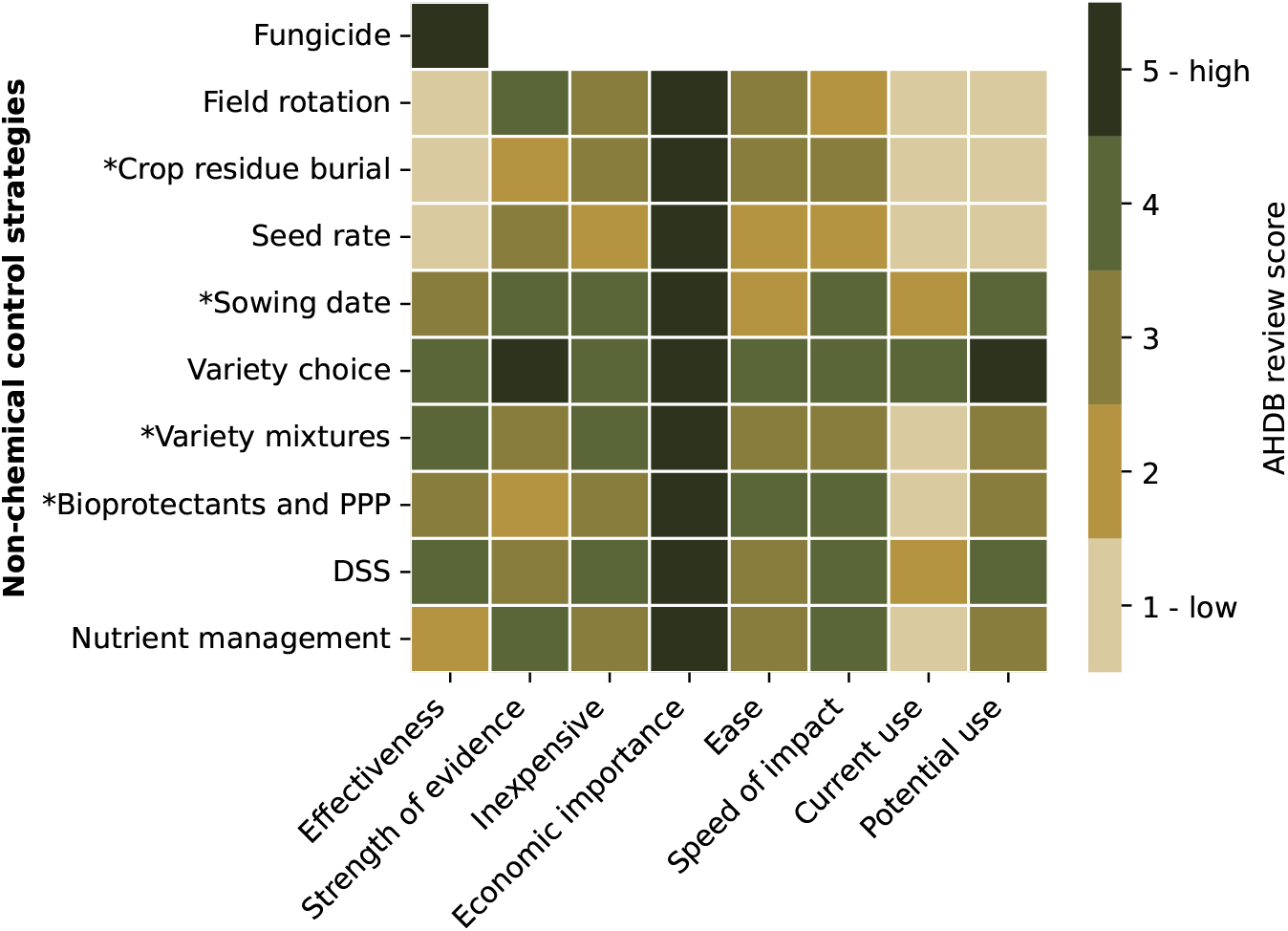
IPM data for Septoria leaf blotch of wheat from the 2021 AHDB Research Review. Blake *et al*. [15] identified and scored IPM measures applicable for this disease by expert opinion on eight qualities, concerning effectiveness of the control, economic factors, and ease/speed of implementation. As in the original review, we display the effectiveness of fungicide for comparison (NB: The other qualities were specific to IPM measures, not fungicide use. As a consequence, there are no data given for these qualities for fungicide in the source). Control measures explicitly included in our study are indicated by a * preceding the control strategy name.

Of the IPM measures identified in the AHDB review, we restricted our examination to those that could be reasonably studied by our mechanistic system model (we expand on the selection of IPM measures for analysis in Section 4). As such, we define the ‘IPM regime’ in this study to be the use of crop residue burial, delayed sowing, variety mixtures, and biocontrols. We implemented each of these IPM measures into the STB model based on a biological understanding of how the control acts and experimental data of their impacts.

#### 2.4.1 Residue burial

Debris and stubble from the previous year’s plants are commonly left in place between seasons. As a consequence, subsequent years’ crops are planted among the residue from previous years. This practice offers certain benefits to growers, such as conserving soil moisture and reducing costs associated with clearing or tilling the field between seasons [15, 22]. However, if infection was present in the previous season, infected tissue can remain on the debris and contribute to the initial infection of the lower canopy in the next season [22–25]. Ensuring that the previous season’s residue is buried before new crops are sown therefore has the potential to reduce the amount of inoculum at the beginning of the season, which could subsequently reduce the size or severity of the outbreak.

Our implementation of residue burial into the model system assumed that this practice would reduce the amount of inoculum initiating STB infection in the lower canopy, thereby reducing the level of initial infection in the *P* class. With Ψ being the base initial value of *P* (Table 1), we assumed that residue burial resulted in an initial value of *pq*Ψ + (1 − *p*)Ψ, where *p* is the proportion of initial infection from debris in the field (rather than from other sources, such as other fields’ debris or crops), and *q* is the contribution to inoculum which remains after debris is buried. Inspecting the two terms in the modified expression for the initial infected leaf area in the lower canopy, the first term *pq*Ψ corresponds to the contribution of initial infection from debris in the field post-residue burial and the second term (1 − *p*)Ψ corresponds to the contribution of initial infection from sources outside the field (assumed to not be impacted by the residue burial intervention implemented in the index field of interest). Brokenshire [23] found that pycnidiospores on unburied debris had a survival rate of 65% after 50 days, while pycnidiospores on buried debris had a survival rate of less than 10% [26]. Based on these empirical data, we considered a range of values for *q* from 0 to 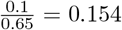 (to 3 significant figures).

The highest reduction in infected leaf area seen in the lower canopy due to debris removal is 75%, seen by Suffert and Sache [25] in the initial stages. However, this reduction is quickly overcome as external sources of infection become more important than debris in contributing to infection in the lower canopy, suggesting that the impact of debris removal could be transient in terms of long-term dynamics [25]. As such, we considered values of *p* that resulted in a disease reduction between 0 and 75%. Each resulting range for *p* depended on the choice of *q* (Fig. S2). The smallest possible range of *p* was (0, 0.775), corresponding to *q* = 0 (when the burial of debris prevents all of its contribution to initial inoculum). The upper limit of this range increased as *q* increased, with the widest possible range for *p* being (0, 0.918) when *q* = 0.154. With no clear choice for the average or ‘most likely’ value of *p*, we chose a default value of *p* = 0.1 (Table 2) based on the strength of evidence that debris removal has a transient impact, and the AHDB review that scored its effectiveness as low (Fig. 2, [15]). We examine in the Discussion the implications of the uncertainty and wide parameter ranges computed here (see Section 4).

**Table 2.**
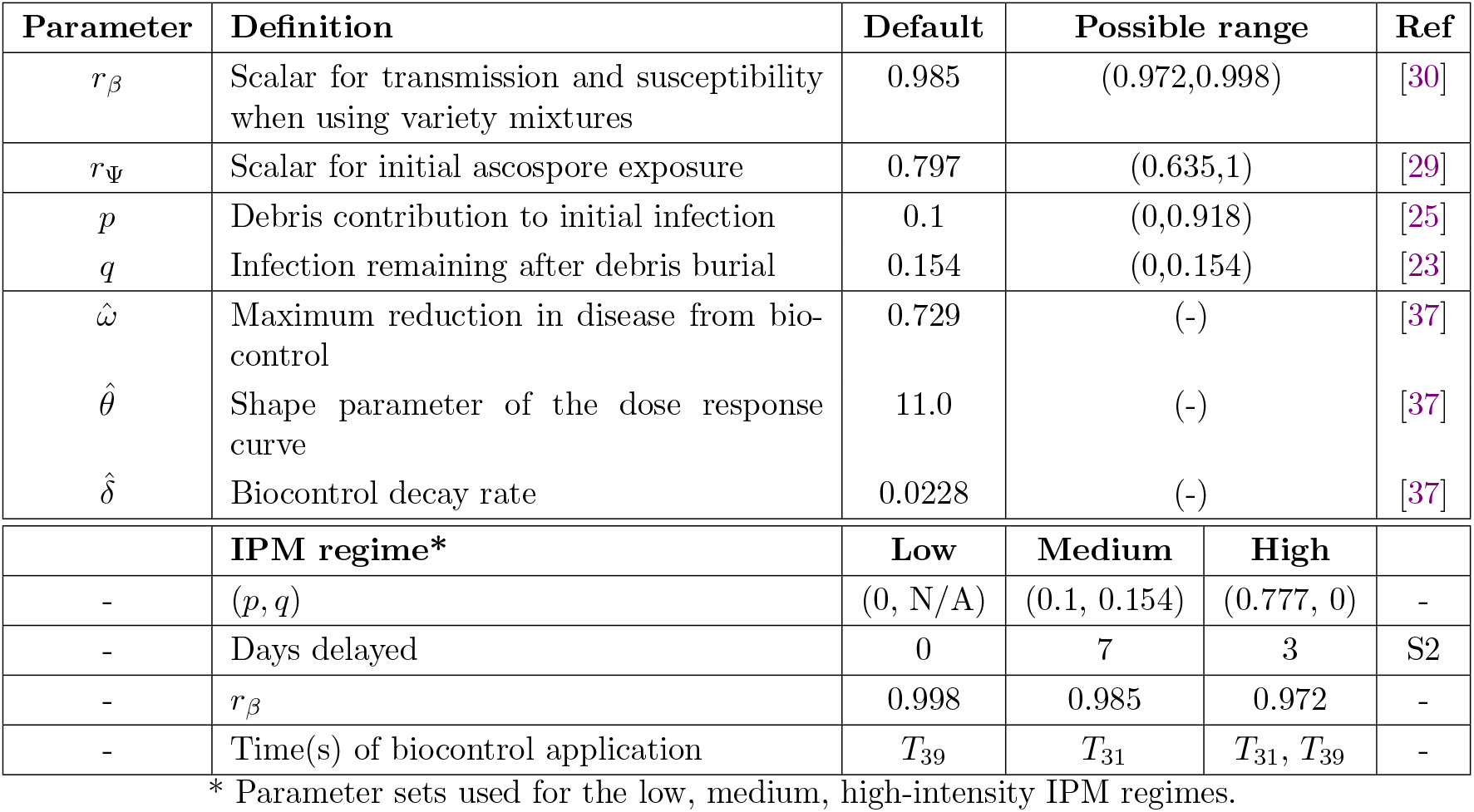
IPM parameter values.

#### 2.4.2 Sowing date

Sowing date has been shown to have an effect on the severity of STB outbreaks [27], with current AHDB guidance suggesting that later sowing increases a crop’s level of potential resistance to STB infection [28]. In many cases, while late sowing results in some level of yield penalty, it also allows the crop to avoid the peak level of inoculum that occurs earlier in the season. As a consequence, the severity of the outbreak can be reduced [28]. We do not account explicitly for reduced crop establishment due to late sowing, which is elaborated on in the Discussion (Section 4).

We implemented late sowing into the model system by introducing a scalar *r*_Ψ_ in [0, 1]. The scalar *r*_Ψ_ allowed us to adjust the magnitude of the initial value of infection in the lower canopy (state *P*), reflecting a reduced exposure to inoculum. We also shifted the initial time of leaf emergence in the model by the appropriate number of degree-days. Late sowing is described in terms of (calendar) days delay after 20th September, matching the assumed sowing date in the Hobbelen *et al*. [9] model.

To compute the reduction in inoculum exposure over time, we fit an exponential decay function to ascospore data collected by Morais *et al*. [29] (Fig. S3). The data estimates the potential amount of ascospores per m^2^ which is produced in a plot with wheat debris. Note that in applying this ascospore decay curve to our model, we do not directly use the calendar dates from the Morais *et al*. [29] experimental data, but shift this data back by 26 calendar days so that the experimental sowing date aligns with the sowing date in our model. Late sown crops will therefore evade the early period of inoculum release, reducing their relative exposure (Fig. S4). We assumed that the level of inoculum exposure *r*_Ψ_ on the first possible sowing date (i.e. 0 days delay) was 1.

We assumed that the maximum number of days a grower would delay sowing by is the latest sowing date such that yield is increased compared to the previous day. In other words, even if sowing an additional day later results in reduced disease prevalence, growers will not do this if the delay to crop growth results in an overall reduction in yield. Under this assumption, the maximum delay a grower would be willing to implement (in the absence of any other controls) is 14 days (corresponding to *r*_Ψ_ = 0.635), as this is the delay that maximises yield. We then assumed that the minimum delay a grower would implement was 0 days, and a median delay of 7 days, which gave values of *r*_Ψ_ = 1 and 0.797 respectively. Under the assumption that growers would like to gain the potential benefits from late sowing, but be reserved due to the risk of uncertain weather conditions when sowing later, we assumed the default value to be the median of a 7-day delay (Table 2).

#### 2.4.3 Variety mixtures

Monoculture farming is a very widely used practice, offering greater uniformity of product for consumers and often simplifying the large-scale growing and harvesting process. However, the genetic uniformity in a varietal monoculture puts crops at greater vulnerability for disease [15, 30, 31]. Increasing genetic diversity within a crop has been shown to reduce both the spread and severity of a disease outbreak [30]; these outcomes are achieved because of a reduction in the probability that infection is successfully transmitted from one plant to its neighbour [31].

Varietal diversity affects both the ability to receive infection (in the *S* state) and transmit infection (from the *I* and *P* state). Existing examples of models of plant disease within variety mixtures include multi-host models that explicitly model the different varieties and have individual parameters for different host-pathogen interactions [32, 33]. We instead sought to implement variety mixtures within our existing transmission model framework in a parsimonious manner that could be parameterised by empirical data. Retaining a single-host type model structure, we assumed a reduction in STB infection that can be achieved under variety mixtures. To further limit parameter dimensionality in our model, we assumed the same value for the scaling parameter acting on susceptibility and transmissibility, *r*_*β*_ in [0, 1]. Incorporating both the scaling on susceptibility and transmissibility resulted in a force of infection term 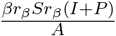(note the repetition of the term *r*_*β*_).

To quantify the reduction in STB infection that can be achieved under variety mixtures, we used the results from Kristoffersen *et al*. [30]. In their meta-analysis of 406 trials that used varietal mixtures (of which the results from 221 trials were ultimately used), they found that the ‘mixing effect’ reduced disease severity by a mean of 9.0%, with lower and upper bounds of 1.0% and 16.3%, compared to the most-grown varieties. For our base model representing an untreated outbreak, we selected the results for untreated trials. Under the assumption that the model presented by Hobbelen *et al*. [9] and the parameters found by Corkley *et al*. [19] represent behaviour of a typically grown variety, we selected the results that compared the varietal mixture outcomes to those of the ‘most-grown varieties’. Given that the use of resistant varieties is already widespread alongside the use of fungicides [15, 34] (Fig. 2), these ‘most-grown varieties’ are typically chosen for their resistant properties. As such, it would be unrealistic to contrast an IPM regime that uses resistant varieties with a fungicide regime that does not use resistant varieties. Accordingly, in our model we implicitly assumed the following two conditions: (i) that varieties planted by the fungicide-using growers were chosen for their resistance, and (ii) variety mixtures used in the IPM regime were a mixture of commonly-used resistant varieties.

We assessed disease severity in our model by integrating between GS69 and GS75, as this is when Kristoffersen *et al*. [30] state that the majority of trials carry out their disease assessment. Fitting *r*_*β*_ to the 1.0%, 9.0% and 16.3% disease reductions resulted in values of 0.998, 0.985 and 0.972, respectively. As the result corresponding to the mean of the original data, we took *r*_*β*_ = 0.985 as our default parameter value for cultivar mixtures (Table 2).

#### 2.4.4 Biocontrols

A number of biofungicide and bioprotectant products are either commercially available, or identified as potentially effective in research studies [35–37]. Many of these products use either bacteria, lower-risk chemical compounds, or other biological agents, to enact either an actively fungicidal effect, or a protectant effect.

These are often applied using the similar methods as chemical fungicides, at similar growth stages, and act through similar mechanisms, though offer a lower level of control. As such, we implemented biocontrol in the model with an analogous form to fungicide (Eqs. (1) and (2)). We used experimental results from 2004 and 2006 reported by Kildea *et al*. [37]. The authors found that *Bacillus megaterium* was capable of inhibiting the development of STB in small-scale field trials. Treatments were applied at GS31 and GS39, to align with conventional fungicide treatment procedures. We used the function

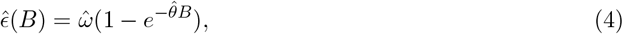

with *Bacillus megaterium* treatment solution concentration

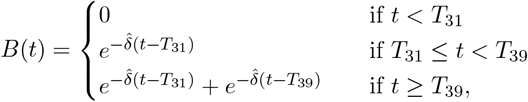

where we fit for the parameters 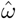(maximum reduction in disease),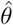 (dose response parameter) and 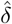 (decay rate) by minimising the distance between the function and the data from Kildea *et al*. [37]. After fitting to both the 2004 and 2006 experimental data, we chose to use the fitting result from the 2004 data, as the 2006 simulation had a visibly worse fit to the no control data (Fig. S5). Our resultant parameter set was 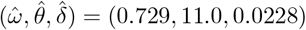. We acknowledge that for this IPM control we have a set of fixed parameter values only, without a potential range. Furthermore, the experimental data only looks at infection prevalence on the flag leaf (the top leaf), while our model would ideally be fit to prevalence in the whole upper canopy. Data limitations prohibited the acquisition of a potential range for these parameters, with further comment on these challenges given within the Discussion (Section 4). As such, unlike the three other control measures, the variation in scenarios explored here is not the range of plausible values, but instead three different application scenarios (as is commonly done experimentally and in practice [35]): one application at *T*_39_, one application at *T*_31_, and one application at both *T*_31_ and *T*_39_. We defined these as the lower bounding, median/default, and upper bounding scenarios respectively (Table 2), based on their comparative outcomes in terms of yield and infection prevalence.

### 2.5 Simulation outline

We produced all model simulation scripts and figures in Python 3.9.12. Code and data for this study are available at https://github.com/emrvincent/IPM_s_tritici_modelling.

#### 2.5.1 Individual IPM measures

To allow us to assess the relative effectiveness of each IPM measure, as well as the potential variability in outcomes between IPM measures and due to parameter uncertainty, we first ran simulations with each IPM measure implemented individually, in an isolated field. We performed model simulations for each of the four IPM measures using the base ODE system (given in Eq. (7)) with modifications relevant to that IPM measure, as described in Sections 2.4.1 to 2.4.4.

For each control measure we output both the infection prevalence over time (defined as the percentage of infectious leaf area; 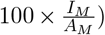 and the relative yield. For comparative purposes, we also generated the same metrics for a ‘no control regime’ and a ‘fungicide regime’. Note that although the no control regime is not typically used in practice, we considered it here for comparative purposes. We implemented the fungicide regime as described in Section 2.3.

#### 2.5.2 Full IPM regime

We next ran simulations to compare three possible field regimes, each employed in an isolated field: the ‘no control regime’, a ‘fungicide regime’ (implemented as described in Section 2.3) and a full ‘IPM regime’. The IPM regime corresponded to the simultaneous implementation of all four of our IPM control measures; residue burial, sowing delay, variety mixtures, and biocontrols.

For each regime we output the infection prevalence over time and the relative yield. The system of ODEs describing the model under the IPM regime is

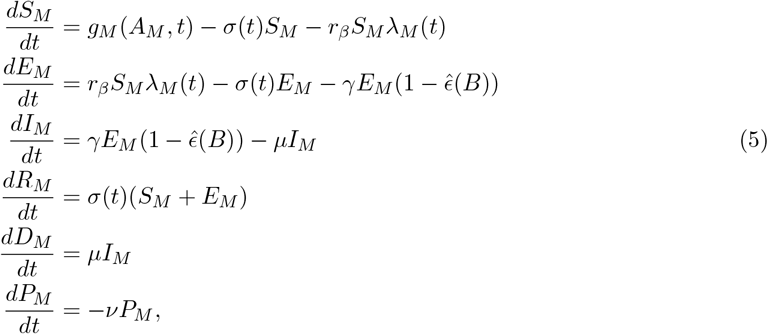

where 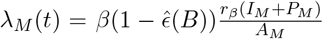. As in the single field fungicide model, *g*_*M*_ is identical to the function *g* in the base system, whilst parameter values and *σ*(*t*) are also the same as in our base system [19] (Table 1 and Section S1). In this case, the initial conditions at the beginning of a year’s growing season were (*S, E, I, R, D, P*) = (0.05, 0, 0, 0, 0, *r*_Ψ_(*pq*Ψ +(1 −*p*)Ψ)), with the initial condition for *P* modified by the control measures. All IPM parameters that were derived in Sections 2.4.1 to 2.4.4 are listed in Table 2.

We ran the IPM regime under three possible scenarios, informed by the range of potential values that our fitting produced for each intervention (selected parameter values for each regime are given in Table 2). Our assumptions for the parameter values selected for each regime intensity were as follows. For variety mixtures, the medium intensity regime used the default value (0.985), and the low and high intensity regimes used the boundary values (0.998 and 0.972, respectively). For delayed sowing, in the low intensity regime we assumed no sowing delay, in the medium intensity regime used the default value of a 7 day delay, and for the high intensity regime we used the value that optimised yield (3 days delay) (Section S2). For residue burial, implementation of the low intensity regime assumed that residue contributed negligibly (*p* = 0) to the initial inoculum, the medium intensity regime used the default values (*p* = 0.1, *q* = 0.154), and the high intensity regime assumed that residual burial contributed the maximal value our fitting allowed (*p* = 0.777, *q* = 0). Finally, the implementation of biocontrol in each regime corresponded directly to the definitions in Section 2.4.4; the low intensity regime had one application at *T*_39_, the medium intensity regime had one application at *T*_31_ and the high intensity regime had both experimental application times.

#### 2.5.3 Multi-field setting

In the multi-field setting, we explored the scenario in which fields are not isolated, but receive infection from all other fields in the system (Fig. 3). In each simulation, we assumed a fixed proportion *ρ* of fields were treated using the IPM regime, while the remaining proportion (1 − *ρ*) of fields were treated using the fungicide regime. Fields of each type were pooled into classes with the subscript *M* and subscript *F* respectively. To model this, we used the system of equations

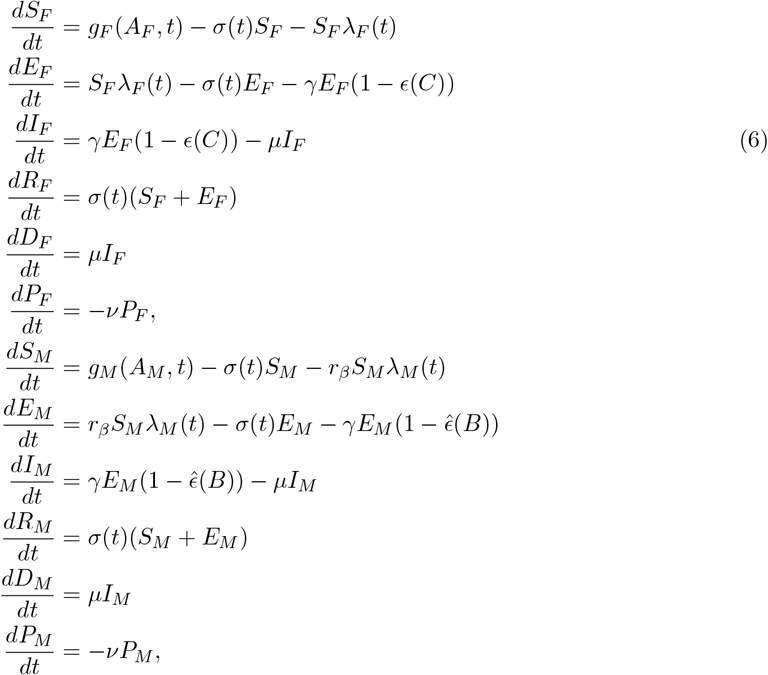

where 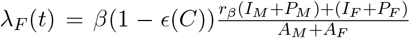 and 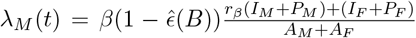. These definitions of the per-capita force of infection on the fungicide and IPM fields respectively account for between-field transmission, with the inclusion of *I*_*F*_ and *P*_*F*_, and *I*_*M*_ and *P*_*M*_ in both transmission terms reflecting the assumption that contact was well-mixed between all fields in the system.

**Fig. 3.**
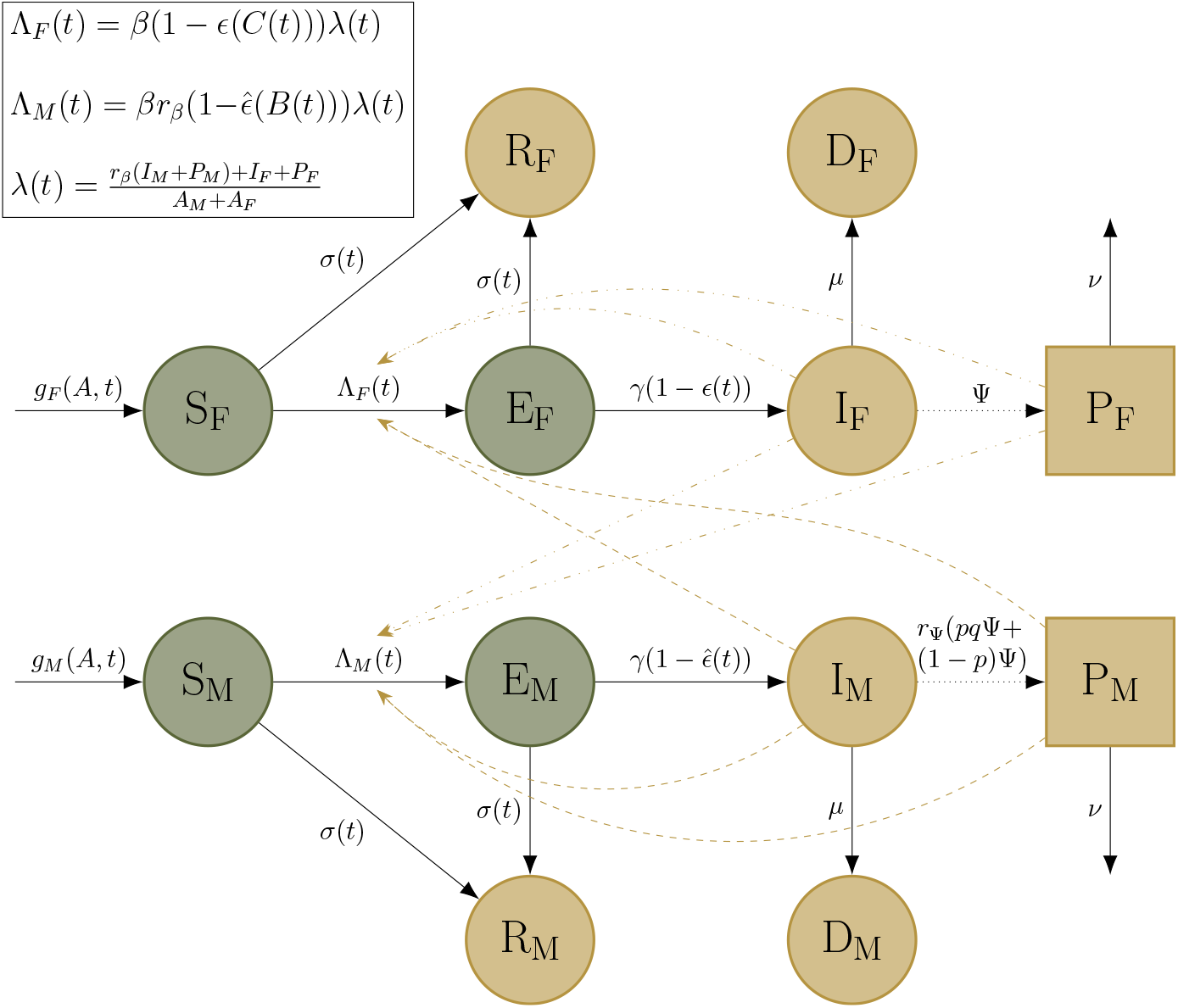
Schematic for the multi-field STB dynamics model. States are divided into fields under the fungicide regime (*F* subscript) and those under the IPM regime (*M* subscript). As a result of the control strategies, compared to the base model the two regimes have different rates on the solid transition arrows and on the dotted between-season arrows. We also introduce transmission between the two field groups.

To account for the proportion of fields of each regime type, the growth terms *g*_*F*_ and *g*_*M*_ must be re-defined so that the maximum leaf area in each field type was scaled by (1 − *ρ*) and *ρ* respectively;

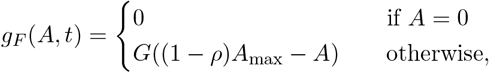

and

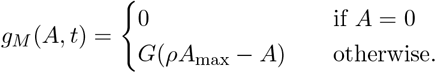

We also applied the same scaling to the initial conditions of each field type, so that the initial condition for this system was

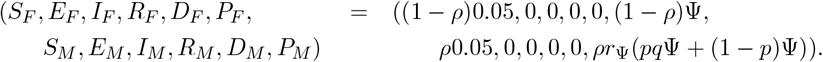

We compared the relative yields of each field type for *ρ* ranging between 0.02 and 0.98, with an increment of 0.02. We output the results for the default IPM parameter set (and under the base values for the fungicide parameters), as well as exploring the system behaviour as we varied the intensity of both regimes (fungicide and IPM). Specifically, we considered all combinations of three different fungicide regime intensities (*ω* = 0.5, *ω* = 0.75, *ω* = 1) and three different IPM regime intensities (low, medium, high), giving a total of nine fungicide-IPM intensity combinations. The three different IPM regime intensities are defined as in the previous, single-field scenario.

#### 2.5.4 Environmental sensitivity and reactive fungicide treatments (DSS)

To account for the real-world variation in environmental conditions, we lastly considered outbreaks of varying severity. We explored this in the single field scenario, implementing this change in outbreak severity by scaling the transmission rate *β* by a constant [9]. We considered three sets of environmental conditions: the standard outbreak severity (*β* scaled by 1), an outbreak that produced infection prevalence aligned with typical high-severity years (*β* scaled by 1.5) [38], and an outbreak that produced infection prevalence aligned with typical low-severity years (*β* scaled by 0.8) [38].

In our main analysis we used our model to consider a strict binary of growers who used fungicide and grower who used IPM without fungicide. Yet, common definitions of IPM regimes do not fully exclude the use of fungicide. Many of them aim to minimise the use of fungicide, but recommend its use in the case of spot treatments, or when a particular disease prevalence is reached. To gain insights on epidemiological and yield outcomes for circumstances where chemical (fungicide) and non-chemical (IPM) measures are used together, we also used our three epidemic severity scenarios to consider the use of reactive fungicide treatment. Real-world implementations of such a strategy are commonly done using Decision Support Systems (DSS). One model used in the IPM Decisions Platform [39] recommends the application of fungicide for STB when a prevalence of 10% is reached. We therefore explored a range of scenarios including that recommendation; the application of fungicide at 10%, 5% and 1% prevalence, along with a scenario that applied fungicide as part of the IPM regime at *T*_32_ by default. In each of these scenarios, we examined the effect of applying different doses of fungicide; a full fungicide application (*ω* = 1), a reduced fungicide application (*ω* = 0.5), and the case of no fungicide treatment (*ω* = 0, equivalent to the IPM regime simulations in Section 2.5.2).

We explored the sensitivity of outcomes to the described environmental and regime changes for all three of our IPM regime scenarios; low-, medium- and high-intensity.

## 3 Results

### 3.1 Individual IPM measures: Sowing date and biocontrol have the greatest impact on yield

In our initial analysis we modelled four different IPM control measures (residue burial, delayed sowing, variety mixtures, biocontrol) applied to an isolated field. We were particularly interested in how their use may impact expected yield and variability in yield (resulting from parameter uncertainty) over a single growing season.

#### Residue burial

When implementing residue burial as the sole control measure, under our default parameter set (*p* = 0.1, *q* = 0.154) it offered little improvement in relative yield or curbing infection compared to having no infection controls (*p* = 0). We observed a relative yield of 0.907 versus 0.900 in the no control scenario (Fig. 4(a)). We observed a peak infection prevalence of 13.2% when using the default parameterisation for residue burial versus a peak infection prevalence of 14.0% in the no control scenario (Fig. 4(b)). Under the best-case parameters (*p* = 0.777, *q* = 0), our model returned a relative yield of 0.973 and the peak infection prevalence was reduced to 4.5%.

**Fig. 4.**
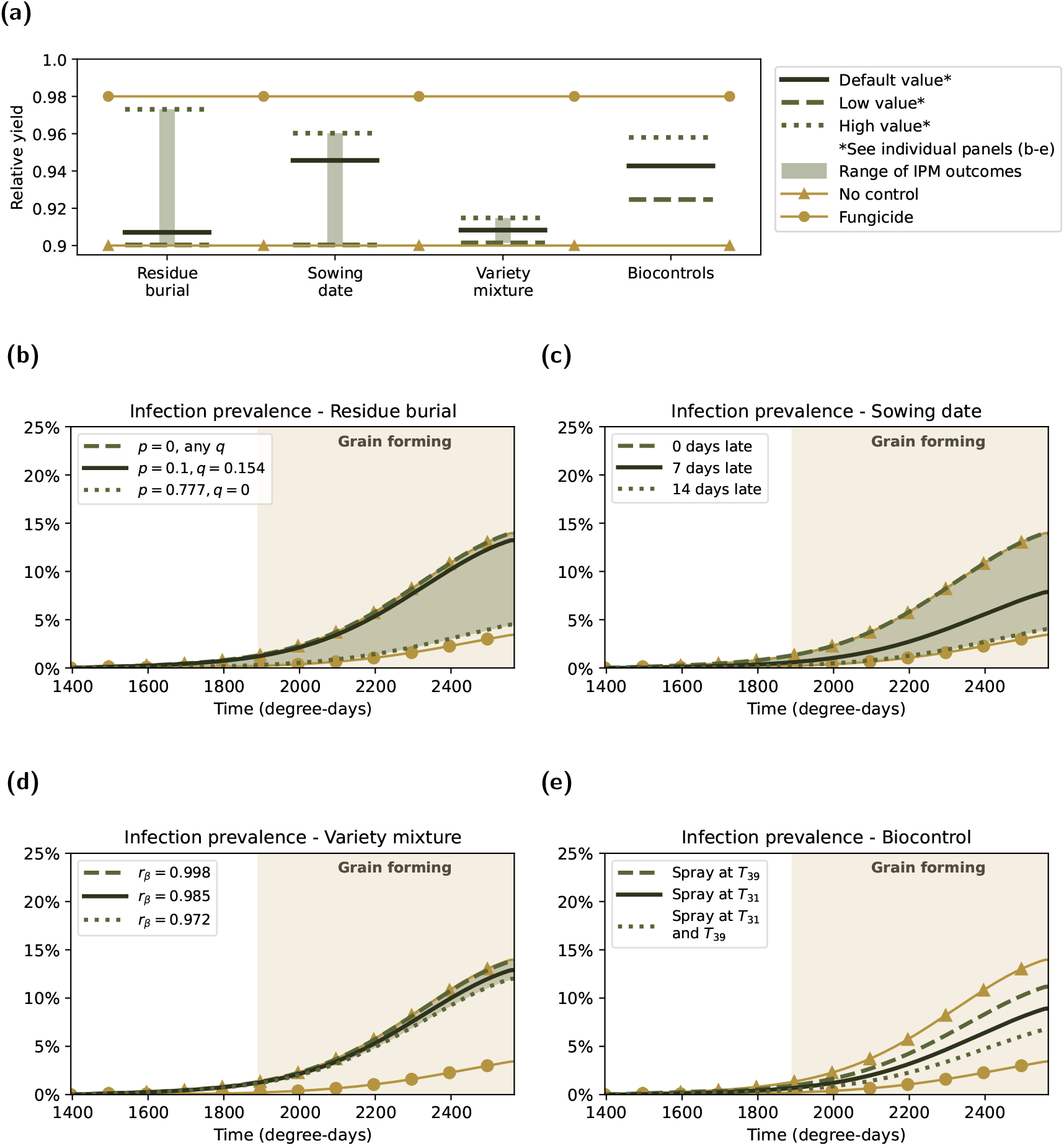
Single season outcomes for each IPM measure implemented individually. For each control, we show the results for the default parameter set by a dark green solid line, the boundary or alternative-implementation results in medium green dashed and dotted lines and, where applicable, the continuous range of possible outcomes the shaded pale green ribbons (we list parameter values and ranges in Table 2). We also show results for the ‘no control’ regime and ‘fungicide’ regime (implemented as described in Section 2.3). Panel **(a)** displays the yield at the end of a season under each of the individual IPM control measures. The legend outside panel **(a)** contains information relevant to all subfigures **(a-e)**, while the legends contained within panels **(b-e)** contain information applicable to each particular IPM measure. Panels **(b-e)** show the infection prevalence (% infectious leaf area,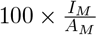) with the following sole IPM control measure applied: **(b)** residual burial; **(c)** delayed sowing; **(d)** variety mixtures; **(e)** biocontrol.

#### Delayed sowing

The second IPM control strategy we considered was delayed sowing. A sowing delay of 14 days resulted in the highest possible relative yield of 0.960, and a peak of infection of 4.2% (Fig. 4(c)). Although sowing delays of more than 14 days result in a lower peak infection prevalence, the yield produced by these scenarios was less than that achieved at 14 days. The lower yield and low is a consequence of the extreme delays to sowing, meaning the plants do not spend enough time experiencing growth before senescence to reach their maximum (healthy) tissue area.

The minimum sowing delay of 0 days (equivalent to the no control scenario) gave the lowest yield from this set of scenarios (0.900). At the intermediate value of a 7-day sowing delay, the relative yield was 0.946 and the peak infection prevalence was 7.9%.

#### Variety mixtures

For the variety mixture control measure, the default parameterisation resulted in a relative yield of 0.908. The best- and worse-case parameterisation increased the relative yield by 0.007 to 0.915, and decreased the relative yield by 0.007 to 0.901 respectively (Fig. 4(a)). Furthermore, the default parameterisation had a peak infection prevalence of 12.9%. The best-case scenario had a 0.9% lower peak prevalence of 12.0%, while the worst-case scenario had a 0.9% higher prevalence at its peak of 13.8% (Fig. 4(d)).

#### Biocontrols

The last IPM control strategy we considered was biocontrol. The functional outputs of Eq. (4) compared to Eq. (2) indicate that biocontrol has a weaker effect in terms of suppressing infection than fungicide (Fig. S6).

We note that since biocontrol could not be parameterised on a range of plausible values (as discussed in Section 2.4.4), this was the only IPM measure in our analysis that had discrete options. A The relative yields for one spray at GS31 and GS39 were 0.943 and 0.925, respectively. The relative yield was higher given two spray treatments at 0.958 (Fig. 4(a)). Applying the biocontrol at only GS31 gave a peak infection prevalence of 8.9%. This contrasted with applying the biocontrol at only GS39, which gave a peak prevalence of 11.2%. A two spray application lowered the peak infection prevalence to 6.8% (Fig. 4(e)).

#### Comparison across individual IPM measures

Our considered range for the variety mixture parameter gave less variability in possible outcomes compared to the variation across the potential scenarios using residue burial or delayed sowing. In the default implementation of each of the four IPM control strategies, sowing date and biocontrol were relatively higher (yields of 0.946 and 0.943) whilst variety mixture and residue burial were relatively lower (yields of 0.908 and 0.907). Across all considered scenarios, we found a maximum relative yield of 0.973 when assuming residue burial was 100% effective at removing inoculum produced on debris and debris being responsible for the majority of the early-season inoculum.

#### Comparison with no-control and fungicide regimes

We initially note that the worst-case implementations of residue burial and sowing date were identical to the no control scenario (Table 2); they therefore produced the same modelled relative yield outcomes. Outside of these two IPM measure parameterisations, all yield values attained by all individual IPM measures were greater than the yield with no control, but less than the yield attained under a fungicide regime (Fig. 4(a)). In terms of infection prevalence, all IPM control scenarios resulted in lower infection prevalence than the no control case, but consistently higher infection prevalence compared to the fungicide control regime (Figs. 4(b) to 4(e)).

### 3.2 Single field setting: Full IPM regime offers comparable control to a fungicide regime

We next considered the implications for infection prevalence and yield for three different control regimes (as if implemented in an isolated field): a ‘no control regime’, a ‘fungicide regime’ (implemented as described in Section 2.3) and an ‘IPM regime’ (with simultaneous implementation of four IPM control measures; residue burial, sowing delay, variety mixtures, and biocontrols). To account for varied grower implementation and variation in parameter results, we also considered three different intensities of IPM regime (Table 2).

For relative yield, the estimate under a fungicide regime of 0.980 was higher than the relative yield estimates using either the low- or medium-intensity IPM regimes (relative yields of 0.925 and 0.972), while both outperformed the no control regime (relative yield of 0.900) (Fig. 5(a)). The high-intensity IPM regime, however, produced a higher yield (0.992) than the fungicide regime.

**Fig. 5.**
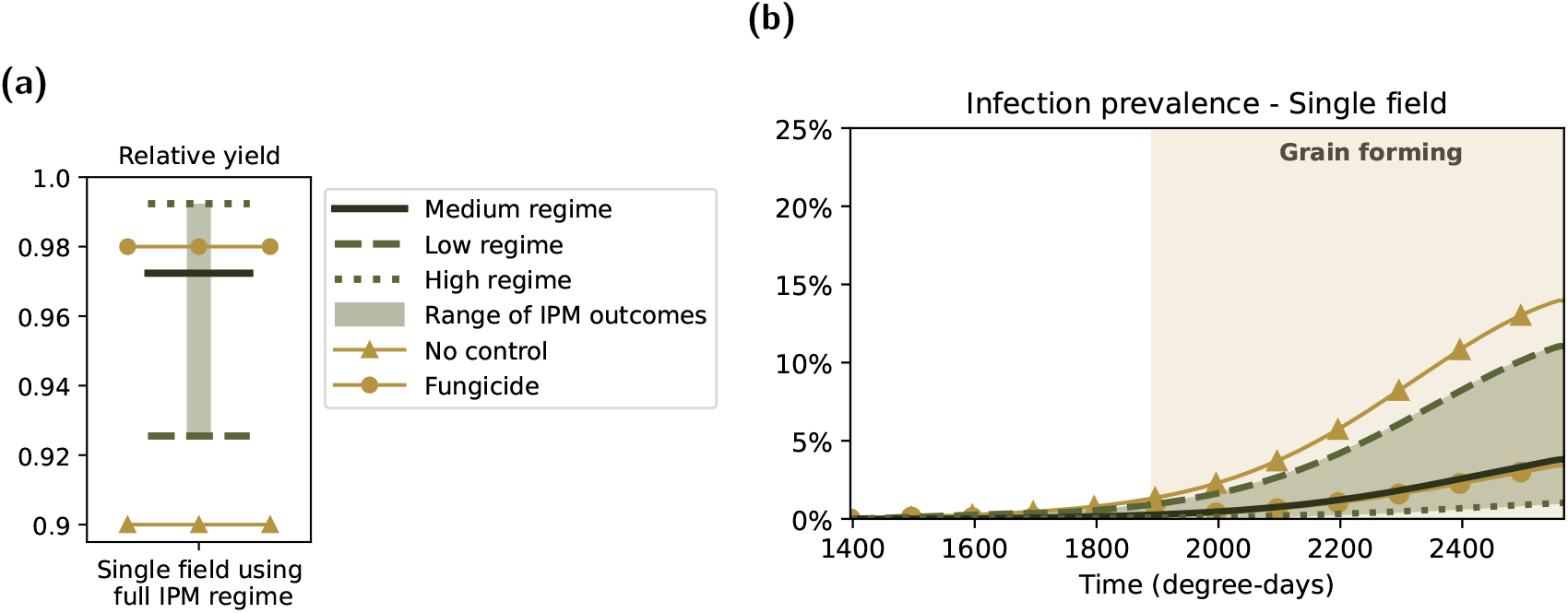
Single season outcomes under three different control regimes. Our three control regimes were ‘no control’, ‘fungicide’, and ‘IPM’ (as described in Section 2.4). **(a)** Relative yield at the end of a season under each regime; **(b)** Infection prevalence (% leaf area infected) over time under each regime.

In terms of infection burden, the medium-intensity IPM regime returned a slightly higher peak infection prevalence (3.81%) than the fungicide regime (3.43%). The low- and high-intensity regimes had peak infection prevalences of 11.1% and 1.0%, respectively. All four control regimes (the three different IPM regimes, and one fungicide regime) outperformed the no control scenario, which had a peak infection prevalence of 14.0% (Figs. 5(b) and S7).

### 3.3 Multi-field setting: Presence of IPM users can improve yield outcomes for all growers

The final component of our analysis involved looking at the outcomes in a system with multiple fields. The multi-field setup allowed for transmission of infection between fields. We examined the effects on the system dynamics of having a varied proportion of fields use an IPM control regime (*ρ*) and the remainder (1 − *ρ*) adopting a fungicide control regime.

We first explore the ‘default’ scenario (medium-intensity IPM regimes and fungicide regimes with *ω* = 1) (top centre panel of Fig. 6). For all values of *ρ*, the IPM regime had a lower relative yield than the fungicide regime. The marginal advantage of fungicide over IPM also increased as *ρ* increased, although the difference between the two never exceeded 0.005.

**Fig. 6.**
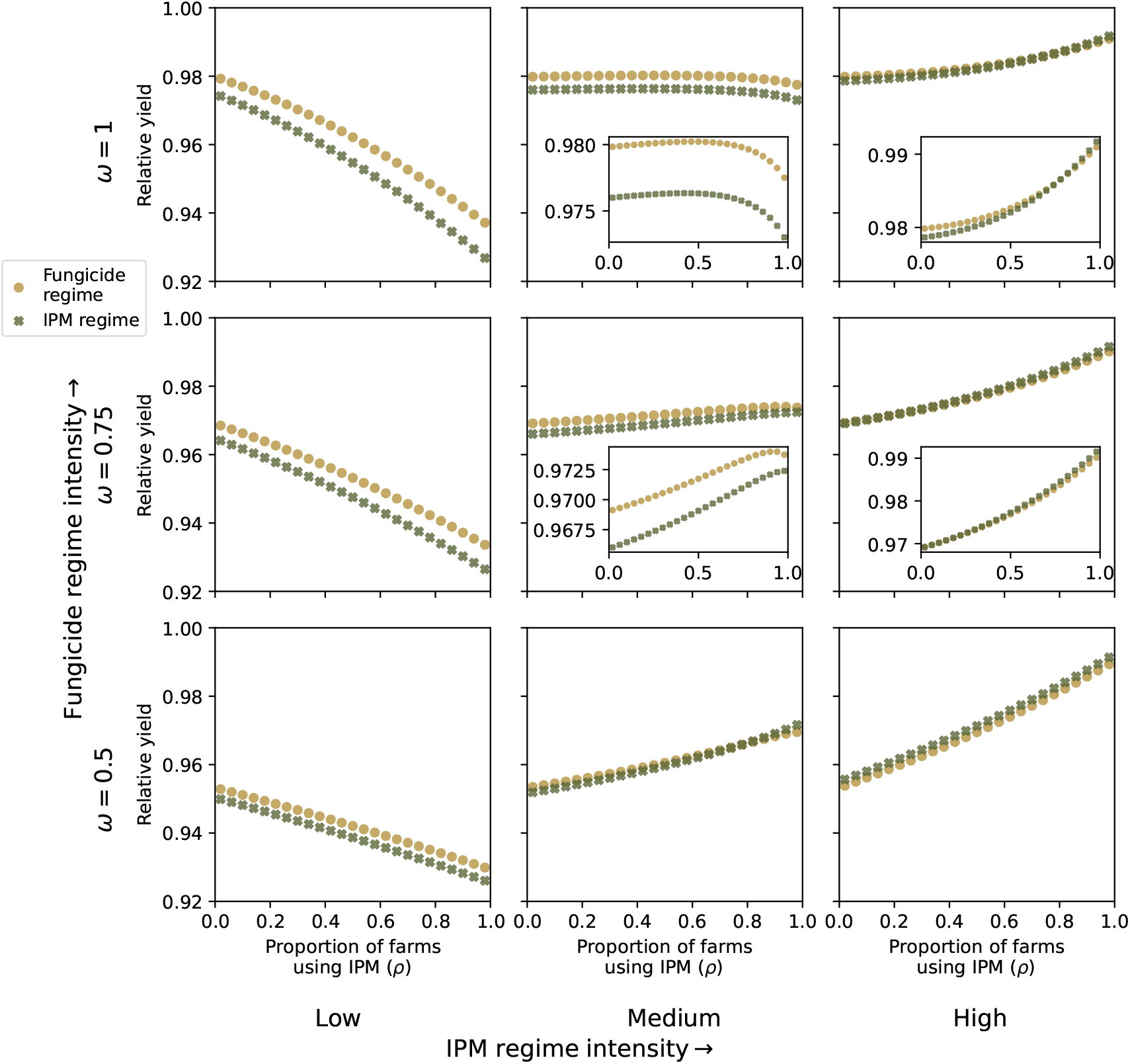
Single season yield outcomes in the multi-field setting. We display end-of-season relative yield for an individual field, within a multi-field system. The parameter *ρ* corresponds to the proportion of fields using the IPM control regime, with the remaining proportion of fields (1 - *ρ*) using a fungicide control regime. We present relative yield outcomes versus *ρ* for three different intensities of IPM control regime and three different intensities of fungicide control regime. Low, medium, and high intensity IPM regimes are defined in Table 2. Low, medium, and high intensity fungicide regimes are defined by *ω* = 0.5, *ω* = 0.75 and *ω* = 1.

At our initial value of *ρ* = 0.02, the IPM and fungicide fields had approximate relative yields of 0.980 and 0.976 respectively. As *ρ* increases from 0.02 to 0.42, increasing the proportion of farms using IPM increased the relative yield for both field types, though this increase is extremely small (below the third significant figure in both groups). In contrast, relative yields decreased from *ρ* = 0.44 onward for the IPM fields and from *ρ* = 0.48 onward for the fungicide fields (Fig. 6). This decrease was more substantial at *ρ* = 0.98 (though we note that this is still a difference of less than 0.01 of relative yield), with the final yields dropping to 0.973 and 0.978 for the IPM and fungicide fields, respectively.

We next inspect the relative yield obtained with different intensities of the two control regimes. At a low level of IPM intensity, and for all three levels of fungicide performance (*ω* = 0.5, 0.75, 1), increasing *ρ* decreased relative yields under either form of control (Fig. 6, left column). However, a higher intensity fungicide regime improved relative yields for both field types, particularly at lower values of *ρ*. Increasing *ω* from 0.5 to 1 gave estimated ranges of relative yield from the IPM fields between 0.926-0.950 (when *ω* = 0.5) and 0.927-0.975 (when *ω* = 1), and estimated ranges of relative yield from the fungicide fields between 0.930-0.953 (when *ω* = 0.5) to 0.937-0.979 (when *ω* = 1).

For the medium-intensity IPM regime, outcomes depended greatly on the fungicide treatment intensity. In most of these scenarios the fungicide fields saw higher relative yields than the IPM fields. The only exception was in the case of *ω* = 0.5 (Fig. 6, middle column, bottom row), where increasing *ρ* resulted in improved relative yields for both field types, with the relative yield for IPM fields surpassing that of fungicide fields at *ρ* = 0.8. In the *ω* = 0.75 scenario (Fig. 6, middle column, middle row), increasing *rho* generally increased relative yield for both field types, with the exception of *ρ* ≥ 0.94 when fungicide fields saw a decline in relative yields. This had a similar drop-off behaviour as the *ω* = 1 scenario (the ‘default’ scenario explored earlier; Fig. 6, middle column, top row), though in that case the decline began at lower values of *ρ*.

With a high IPM regime intensity, relative yields increased as *ρ* increased (Fig. 6, right column). However, as fungicide intensity (*ω*) increased, the outcomes changed from IPM fields consistently outperforming fungicide fields (*ω* = 0.5, Fig. 6, right column, bottom panel), to almost identical relative yields for both farm types (*ω* = 0.75, Fig. 6, right column, middle panel), to higher relative yields in the fungicide fields for all values of *ρ* ≤ 0.76.

### 3.4 Environmental sensitivity and reactive fungicide treatments (DSS): Reactive use of fungicide offers little improvement in high-effectiveness IPM regimes, even under high-severity outbreaks

We explored the environmental sensitivity of outcomes by considering simulations with three different outbreak severities; scaling *β* by 0.8, 1 and 1.5. In these simulations we also explored the controlled use of fungicide alongside an IPM regime, with fungicide being applied when certain conditions are met: 10%, 5% and 1% infection prevalence, and application at *T*_32_.

As expected, more severe outbreaks result in worse relative yield outcomes overall, with values ranging from 0.797-0.993 (Fig. S8). Less severe outbreaks returned better relative yield outcomes, with values ranging from 0.960-0.997. The early application of fungicide (at a lower prevalence threshold) always produced higher estimated relative yields than a later application of fungicide (at a higher prevalence threshold). Under all three outbreak severities, having fungicide applied at *T*_32_ gave the highest relative yield outcomes and lowest peak prevalence. Fungicide application at 10% gave the lowest overall relative yields and highest peak prevalences.

In a low-severity outbreak scenario, some of the fungicide application thresholds were not met (10% prevalence was never achieved, while 5% prevalence was never achieved for the medium-intensity IPM regime, and 1% prevalence was never achieved for the high-intensity IPM regime) (Fig. S8). However, even in the cases where the fungicide application threshold was met and fungicide was applied, the application of greater quantities of fungicide (*ω* = 1 compared to *ω* = 0, 0.5) resulted in little improvement in relative yield. By contrast, in the high-severity outbreak the fungicide application thresholds were often met. When the fungicide treatment was applied it led to substantial improvements in relative yield. For all three outbreak severity conditions, we found the largest improvement in relative yield gained by the use of reactive fungicide treatment under the low-intensity IPM regime and with application of fungicide at *T*_32_. In the low-severity outbreak this maximal improvement corresponded to an increase from a relative yield of 0.960 when *ω* = 0, to a relative yield of 0.982 when *ω* = 1. In the high-severity outbreak this maximal improvement in relative yield was an increase from 0.797 to 0.9043 (Fig. S8).

## 4 Discussion

### 4.1 Findings

With greater constraints on the use of fungicide, the effectiveness of IPM as a control for arable diseases has become increasingly relevant. Previous work has identified IPM measures that are effective at controlling STB and explored their implementation in practice [27, 35, 37, 40]. By incorporating these existing data into an epidemiological model, we explored the potential outcomes of these IPM measures on the overall disease system. We found that individual IPM measures offered differing levels of control, but that an ‘IPM regime’ consisting of the simultaneous implementation of multiple methods offered comparable disease control and yield to a standard fungicide regime. Furthermore, when IPM was implemented at a reasonable level of effectiveness (default/medium-intensity or better), or when fungicide was below its optimal effectiveness (*ω* ≤ 0.75) the presence of IPM-using fields benefitted all growers in a multi-field system. An additional finding was that, in general, improved data on different IPM measures is critical to the ability to judge its effectiveness as a regime, and which measures to deploy concurrently.

For the use of single IPM controls, there was a striking difference between the potentially high effectiveness of sowing date and biocontrols, and the potentially lower effectiveness of residue burial and variety mixtures. Our results suggest that when implemented independently of other IPM measures, delayed sowing has the potential to reduce the severity of STB outbreaks, with similar levels of control offered by biocontrols. These findings are corroborated by the AHDB review (Fig. 2, [15]) that assigned both of these measures an effectiveness score of 3/5 (with 4/5 being the maximum effectiveness score given to any IPM control of STB). In addition, while there is no field data to corroborate these results for biocontrol, our findings for delayed sowing corroborate experimental studies [41]. On the other hand, whereas the study by Bekele and Kassaye [41] found that the decrease in disease severity had no effect on yield, our modelling results found an increased yield with decreasing disease severity.

Our model findings for delayed sowing and biocontrol contrast with those for the implementation of residue burial and variety mixtures, which offered little control over the size of the final outbreak. This finding for variety mixtures is in contrast to the AHDB review (Fig. 2, [15]), which assigns variety mixtures the maximum effectiveness given to a non-fungicide control measure of 4/5. We note that variety mixtures had the most extensive source of parameterisation data, meaning the parameter value considered was the narrowest of the four individual control measures [30]. However, our findings for residue burial are in agreement with the findings of Morais *et al*. [29], Suffert and Sache [25] and the AHDB review data (Fig. 2, [15]). Residue burial providing little control on the amount of infection during an outbreak appears to result from the fact that even when the burial of debris is very effective at eliminating inoculum from this source (*q* = 0.154 in our default parameter set), larger-scale sources of primary inoculum are likely to play a bigger role in initiating an outbreak [25]. Furthermore, the amount of local initial inoculum is unlikely to meaningfully limit infection development, as the presence of larger-scale sources of inoculum during the emergence of the first leaf mean that its infection is likely to occur almost immediately upon its emergence, regardless of the presence of early within-field inoculum [29]. It is important to note that the effectiveness of residue removal varies a lot depending on how important debris is as a contributor of inoculum. Although its contribution is believed to be low [25, 29, 42], it is known to be extant [23, 43], and has not been well-quantified. We therefore encourage the collection of additional empirical data to help quantify this process.

In the single field setting, we found that IPM was able to serve as an effective measure of control against STB, in agreement with previous theoretical and experimental findings about STB [30, 41, 44] and other diseases [44–46].

When considering the multi-field setting, we found that increasing the presence of IPM regimes within the system can offer increased yields to all growers. This was the case if the IPM regimes are sufficiently effective; that is, our definitions of medium or high intensity (Fig. 6). We hypothesise these yield benefits arise when more growers use IPM control because, while the fungicide regime offers protection only to the target field, the IPM regime offers control to other fields in the system by several mechanisms: (i) reducing the force of infection it emits due to the use of variety mixtures (*r*_*β*_); (ii) reducing the initial infection due to debris removal; (iii) reduced overall ascospore exposure, and; (iv) the asynchronous timing of initial infections between the two farm types.

We also identified circumstances (the medium-intensity IPM regime, with *ω* = 0.75 or 1) where if the proportion of fields using IPM was already at a high level, then further increasing the proportion of farms who use IPM could result in a decrease in yield. This appears to occur due to the fact that under these circumstances, the proportion of farms using fungicide becomes too small to realise the benefits that the fungicide control provides; that is, to reduce infection in those fields, which occurs just prior to the emergence of the IPM fields. In other words, having few farms using fungicide in the scenarios where fungicide is very effective by comparison to IPM reduces the possible benefit this control offers to the system, while also disrupting the ‘asynchronicity’ effect, as the majority of infection emerges with the delayed-sown IPM fields.

Our modelling approach is similar to that taken in several other studies in considering a mechanistic, S-I system of equations where multiple strategies are present, and infection is spread between all fields in the system [18, 47]. Although we were not able to identify any studies that have previously implemented this type of method to compare an IPM-only strategy with a fungicide-only strategy, Murray-Watson and Cunniffe [18] investigated the choice between resistant and tolerant varieties, McQuaid *et al*. [47] considered the use of clean planting material in the growing of cassava, and Radici *et al*. [48] considered the optimisation of fungicide deployment to reduce their use to prioritised scenarios, without compromising disease control. All of these would be considered to be IPM measures.

Murray-Watson and Cunniffe [18] found that increasing the proportion of growers using resistant varieties decreased the infection level in both users and non-users of the control. We observed a similar phenomenon in our system; an increasing proportion of IPM growers offers increased benefits to other farmers in the system, in addition to the users themselves. However, the opposite trend was present in their results for the use of tolerant varieties; in this case, the increased presence of tolerant varieties resulted in higher overall infection presence for control users. This was because tolerant varieties still harbour the pathogen, in contrast to resistant varieties which do not.

The previously mentioned studies focus primarily on grower behaviour dynamics in the system. We were able to identify few similar studies that have studied purely epidemiological modelling outcomes in the context of IPM. The system on which our model was based has been used to investigate the epidemiological outcomes of different control strategies for STB. While much of this work investigates the development of resistant strains of infection under various regimes of fungicide control [9–12], one study by Taylor and Cunniffe [14] does consider the implementation of an IPM strategy. They found that disease resistant cultivars were able to decrease infection and increase yield, and we found improvements in the same outputs under the use of an IPM regime; although our IPM regime did not include resistant varieties, the IPM measure considered in this study. Nonetheless, our approach differs to theirs in that we found IPM to be an effective control in the absence of fungicide, while Taylor and Cunniffe [14] showed that an IPM measure can be used to reduce the quantity of fungicide which is needed for control. Other notable studies that take a modelling approach to the use of IPM are those by Tang *et. al* [45, 49, 50]. Similarly to our work, they found that sufficiently effective or well-implemented IPM regimes were capable of effectively suppressing pest population. These studies differ from ours in methodology in a number of ways however: in their consideration of pests and ours of infectious disease; their approach being that of dynamical system analysis and ours of numerical solutions; and theirs of considering an IPM regime which includes the use of chemical pesticide, while we primarily consider the use of IPM and fungicide in the absence of one another.

### 4.2 Limitations

The model we have presented is necessarily a simplified representation of reality. It is therefore important that we consider the modelling assumptions made and their potential limitations. We elaborate here on four items. These are the implications of considering IPM and fungicide to be two completely distinct regimes, the exclusion of certain IPM measures from our modelling framework, the potential compounding effect of concurrent implementation of multiple IPM controls, and the variability in IPM parameter ranges used in our model construction.

First, to make the comparison between the fungicide and IPM regimes more straightforward, we excluded any use of fungicide from our IPM regime, and vice versa. Although we recognise this as a simplifying assumption, it allowed us to more clearly contrast different IPM options with a common control method that did not use IPM. We did however explore the use of reactive fungicide treatment, as employed in Decision Support Systems (DSS) in Section 3.4. To further limit the overlap between these two regimes, as addressed in Section 2.4.3, we also omit the explicit consideration of resistant varieties.

Second, we chose to include only a subset of the IPM measures identified in the AHDB review (Fig. 2, [15]) as having the potential to control STB of wheat, in our model. This was done due to the unsuitability of including certain measures in our model. The omitted IPM measures were field rotation, seed rate, and nutrient management. We omitted field rotation because our model contains a population-average for growers using IPM or fungicide, rather than characterising individual fields that would be better suited for studying field rotation. We omitted seed rate and nutrient management as these measures would affect plant population/leaf density in a way that cannot be readily implemented into our current model. Given that each of these measures (field rotation, seed rate, and nutrient management) have been identified as reducing overall infection in the system (at varying levels of effectiveness), it seems likely that were these additional control methods included in the system, they would offer at least as much suppression of infection as our present model.

For our implementation of delayed sowing we also did not explicitly account for a reduced yield due to failures in establishment. Although it is acknowledged that later sowing reduces the rate of successful seed establishment, this is typically accounted for by increasing the seed rate sown [20, 51, 52]. That being said, delayed sowing can still implicitly result in yield reductions within our model. Based on the date the crop is sown, some degree-day penalty is incurred to upper canopy emergence, ultimately reducing the growth period of the plant before senescence (Fig. S4).

Third, we did not have sufficient data to account for any compounding effect that these IPM measures would have when acting in combination. Nonetheless, we made subjective considerations for the implementation of each IPM measure when used in conjunction with other IPM measures. For example, we assumed that when sowing was delayed, the time at which biocontrol was applied shifted by the corresponding number of degree days. We also assumed that the reduction in inoculum due to sowing delay would act on both the *pq*Ψ component, and the (1 − *p*)Ψ component of the initial condition. In other words, we assumed that late sowing would affect exposure to inoculum from both debris and other sources.

Lastly, we note that there is a high amount of variability in the certainty of some IPM parameters in our model construction. In the case of debris burial, although the relative contribution of local debris to the initial pool of inoculum has been qualitatively noted [25, 26], it has not been quantified in a way that allowed it to be integrated into our model. Suffert and Sache [25] do numerically compute the reduction in disease severity early in an outbreak, attributable to chopping or removing debris. However, we deemed these data as not suitable for our purposes as these methods of debris control may have different impacts compared to burial. Additionally, the experimental data in Suffert and Sache [25] was taken only in the lower canopy and at timepoints outside of our model timeframe. We encourage the collection of further data to improve our understanding of the relative contributions of sources of inoculum over time during the initial stages of an STB outbreak. In the case of biocontrols, there was almost no experimental literature that we could identify that quantified the reduction in disease severity due to the use of biocontrol on STB on a field-scale, at multiple timepoints, although many have done this in-vitro [44, 53], or in-vivo on an individual plant level [44, 54, 55], at a small number of timepoints. Perelló *et al*. [36] and Mann *et al*. [35] are the only studies we have identified at the time of writing that have quantified the field-level impact of a biocontrol on STB severity. These studies are themselves limited as both assessed disease severity at only one timepoint, which made them unsuitable for fitting the three necessary parameters in our model. To accurately predict their ability to control disease when employed by growers in practice, we advocate for the need for collection of experimental data on the effectiveness of biocontrol measures on a field-scale.

We also acknowledge that the scope of our study has focused on biological and epidemiological outcomes. We recognise that reducing the outcomes only to yield excludes several key factors that can affect growers’ decisions in practice. Though a more extensive consideration of human and market factors is beyond the scope of this study, exploration of these factors warrants attention in future work. Such factors include the cost of the IPM and fungicide control strategies (both direct and indirect), and the market price of the different yields (non-fungicide treated wheat may be sold as organic and thus for a higher price).

### 4.3 Further work

In the present work we considered a purely epidemiological model of the disease system. We have yet to account for any social or behavioural factors that would influence the choice of control adopted. Across the literature, it is acknowledged that profitability (of which one may consider cost, yield, time, and ease of implementation) is a major factor in growers’ choice of control using IPM, or other methods [15, 56, 57]. The AHDB review data used in the present study (Fig. 2, [15]) also acknowledges the varying ease of implementation and cost that characterises different IPM measures. Deepening our understanding of adoption of plant disease controls and grower behaviour towards disease management is an inherently interdisciplinary challenge. There is consequently an urgent need to pursue interdisciplinary approaches to understand the epidemiological and behavioural factors behind such decision making.

In light of this, future work will focus on combining this epidemiological model with a behavioural model for growers intentions towards disease control. Development of a conjoined epidemiological-behavioural model will enable us to extend the analysis beyond a single growing season, as the initial conditions for subsequent seasons will be defined by both infection prevalence and the perceived success of each control strategy by growers in the previous season. This work would be in line with previous literature considering the interactions between grower behaviour and the control of a disease system [17, 18, 47].

Moreover, considering scenarios over multiple growing seasons will enable assessment of the potential implications on estimated yield under the different IPM measures of other important processes occurring over multiple years (such as pathogen resistance to chemical interventions and climate change).

For many crops, chemical control is likely to be a necessary part of crop protection for the foreseeable future [58]. However, the intensive use of chemical control measures has led to challenges in the resistance of pathogens to chemical controls. The implementation of resistance management strategies has become paramount in prolonging the effectiveness and usefulness of chemical controls to growers [59, 60]. IPM can contribute to chemical control resistance management by limiting the exposure time of the pathogen population to the chemical treatment and by keeping to a minimum the chemical treatment input required for disease control. For example, experimental observations are corroborated by modelling studies suggesting that cultural control measures that delay inoculum arrival can contribute to resistance management [61, 62]. With our model framework, considering time horizons spanning multiple growing seasons offers scope for explicitly modelling how the joint use of chemical controls and multiple IPM measures can impact the evolutionary processes associated with *Z. tritici* resistance to chemical controls.

With sowing date decisions having the biggest impact on estimated yield in our single growing season scenarios, a notable area for further analysis will be the implications of climate change on the trade-off between late sowing to avoid disease and early sowing to maximise yield. Changes in climate are likely to affect the timing of certain life stages, which would in turn affect considerations around optimal sowing date [63]. High temperatures and drought are likely to be two key climactic factors that would affect the development of wheat and the transmission of pathogens [64]. Gvozdenac *et al*. [65] note that climate change is likely to increase the pathogen load of diseases that are residue-borne; shifts in residue-borne pathogen load may affect the control strategy of residue burial, either by increasing its importance or reducing its effectiveness. Those authors also summarise several impacts of climate change on other foliar diseases of wheat, which may also prove to be applicable to STB, including an expanded overwintering area of wheat stem rust. Finally, although DSS is not considered as a part of the core IPM regimes in the present study, it is noted by Zayan [66] that DSS, including forecasting and early warning systems, are likely to become more crucial for growers and decision-makers as the frequency of climate phenomena increases.

### 4.4 Conclusion

In this study, we have parameterised a novel model of STB transmission, fungicide control and IPM measures to investigate the potential impact on yield of different control strategies. We found that the presence of fields using IPM may offer indirect control to all growers in the system. We also identified where detailed empirical studies will help reduce uncertainty in model parameterisation and improve the robustness of model projections; specifically, quantifying contribution of crop debris to inoculum and the collection of experimental data on the effectiveness of biocontrol measures on a field-scale. Overall, the results suggest that wheat growers can consider IPM as a viable alternative to a conventional fungicide regime whilst maintaining yield, particularly as the use of chemical fungicides comes under increasingly tighter constraints. As growers not participating in IPM control could indirectly benefit from its presence, this has interesting implications for incentivisation of IPM uptake, as we see a free-rider effect that has been observed in other cases of crop disease management [16–18].

## Author contributions

**Elliot M.R. Vincent:** Data curation, Formal analysis, Methodology, Software, Validation, Visualisation, Writing - Original Draft, Writing - Review & Editing.

**Edward M. Hill:** Conceptualisation, Methodology, Supervision, Validation, Visualisation, Writing - Original Draft, Writing - Review & Editing.

**Stephen Parnell:** Conceptualisation, Methodology, Supervision, Validation, Visualisation, Writing - Original Draft, Writing - Review & Editing.

## Data availability

All data utilised in this study are publicly available, with relevant references and data repositories stated within the main manuscript. The code repository for the study is available at: https://github.com/emrvincent/IPM_s_tritici_modelling. Archived code at time of publication: https://doi.org/10.5281/zenodo.15498417.

## Code availability

The code repository for the study is available at: https://github.com/emrvincent/IPM_s_tritici_modelling. Archived code: https://doi.org/10.5281/zenodo.15498417.

## Acknowledgements

We thank Peter Hobbelen for sharing the time data from *Delaying Selection for Fungicide Insensitivity by Mixing Fungicides at a Low and High Risk of Resistance Development: A Modeling Analysis* [9], and Robert Lillywhite for his input about current biocontrol availability.

## Supporting Information

**S1 Text. Supporting information for ‘Modelling the effectiveness of Integrated Pest Management strategies for the control of Septoria tritici blotch’**.

This supplement consists of the following parts: (1) Base model; (2) Variation in optimal sowing date; (3) Additional figures.

## References

[1] Van Dijk M, Morley T, Rau ML, Saghai Y. A meta-analysis of projected global food demand and population at risk of hunger for the period 2010–2050. Nature Food 2(7):494–501 (2021).

[2] Department for Environment, Food & Rural Affairs. United Kingdom Food Security Report 2021: Theme 1: Global Food Availability (2021). URL https://www.gov.uk/government/statistics/united-kingdom-food-security-report-2021/united-kingdom-food-security-report-2021-theme-1-global-food-availability. [Online] (Accessed: 22 May 2025).

[3] Agriculture and Horticulture Development Board. Septoria tritici in winter wheat (2024). URL https://ahdb.org.uk/knowledge-library/septoria-tritici-in-winter-wheat. [Online] (Accessed: 22 May 2025).

[4] European Court of Auditors. Sustainable use of plant protection products: limited progress in measuring and reducing risks (2020). URL https://www.eca.europa.eu/Lists/ECADocuments/SR20_05/SR_Pesticides_EN.pdf. [Online] (Accessed: 22 May 2025).

[5] Ridley L, Parrish G, Chantry T, Richmond A, MacArthur R, et al. Arable crops in the United Kingdom 2022 (2024). URL https://pusstats.fera.co.uk/api/report-download/699. [Online] (Accessed: 22 May 2025).

[6] Kogan M. Integrated pest management: historical perspectives and contemporary developments. Annual review of entomology 43(1):243–270 (1998). doi:10.1146/annurev.ento.43.1.243.

[7] Collier R. Pest insect management in vegetable crops grown outdoors in northern Europe– approaches at the bottom of the IPM pyramid. Frontiers in Horticulture 2:1159375 (2023). doi:10.3389/fhort.2023.1159375.

[8] Adamson H, Turner C, Cook E, Creissen HE, Evans A, et al. Review of evidence on Integrated Pest Management. DEFRA. (2020).

[9] Hobbelen P, Paveley N, Van den Bosch F. Delaying selection for fungicide insensitivity by mixing fungicides at a low and high risk of resistance development: A modeling analysis. Phytopathology 101(10):1224–1233 (2011).

[10] Hobbelen P, Paveley N, Oliver R, Van den Bosch F. The usefulness of fungicide mixtures and alternation for delaying the selection for resistance in populations of Mycosphaerella graminicola on winter wheat: a modeling analysis. Phytopathology 103(7):690–707 (2013).

[11] Elderfield JA, Lopez-Ruiz FJ, van den Bosch F, Cunniffe NJ. Using epidemiological principles to explain fungicide resistance management tactics: Why do mixtures outperform alternations? Phytopathology 108(7):803–817 (2018).

[12] Taylor NP, Cunniffe NJ. Optimal resistance management for mixtures of high-risk fungicides: robustness to the initial frequency of resistance and pathogen sexual reproduction. Phytopathology 113(1):55–69 (2023).

[13] Van den Berg F, Van den Bosch F, Paveley N. Optimal fungicide application timings for disease control are also an effective anti-resistance strategy: a case study for Zymoseptoria tritici (Mycosphaerella graminicola) on wheat. Phytopathology 103(12):1209–1219 (2013).

[14] Taylor NP, Cunniffe NJ. Modelling quantitative fungicide resistance and breakdown of resistant cultivars: designing integrated disease management strategies for Septoria of winter wheat. PLOS Computational Biology 19(3):e1010969 (2023).

[15] Blake, Cook, Godfrey, Tatnell, White, et al. Research Review No. 98 - Enabling the uptake of integrated pest management (IPM) in UK arable rotations (a review of the evidence). Technical report, AHDB (2021).

[16] Murray-Watson RE, Cunniffe NJ. Expanding growers’ choice of plant disease management options can promote suboptimal social outcomes. Plant Pathology 72(5):933–950 (2023).

[17] Milne AE, Bell JR, Hutchison WD, van den Bosch F, Mitchell PD, et al. The effect of farmers’ decisions on pest control with Bt crops: a billion dollar game of strategy. PLoS computational biology 11(12):e1004483 (2015).

[18] Murray-Watson RE, Cunniffe NJ. How the epidemiology of disease-resistant and disease-tolerant varieties affects grower behaviour. Journal of the Royal Society Interface 19(195):20220517 (2022).

[19] Corkley I, Mikaberidze A, Paveley N, van den Bosch F, Shaw MW, et al. Dose Splitting Increases Selection for Both Target-Site and Non-Target-Site Fungicide Resistance—A Modelling Analysis. Plant Pathology (2025).

[20] HGCA. The wheat growth guide, Spring 2008, second edition (2008). URL https://farmpep.net/sites/default/files/2022-12/Wheat%20Growth%20Guide_2008_UK%20%281%29.pdf. [Online] (Accessed: 22 May 2025).

[21] AHDB. Fungicide performance in cereals and oilseed rape (2025). URL https://ahdb.org.uk/knowledge-library/fungicide-performance-in-cereals-and-oilseed-rape. [Online] (Accessed: 22 May 2025).

[22] Bailey K, Duczek L. Managing cereal diseases under reduced tillage. Canadian Journal of Plant Pathology 18(2):159–167 (1996).

[23] Brokenshire T. Wheat debris as an inoculum source for seedling infection by Septoria tritici. Plant Pathology 24(4):202–207 (1975).

[24] Scott P, Sanderson F, Benedikz P. Occurrence of Mycosphaerella graminicola, teleomorph of Septoria tritici, on wheat debris in the UK. Plant Pathology 37(2):285–290 (1988).

[25] Suffert F, Sache I. Relative importance of different types of inoculum to the establishment of Mycosphaerella graminicola in wheat crops in north-west Europe. Plant Pathology 60(5):878–889 (2011).

[26] Suffert F, Sache I, Lannou C. Early stages of septoria tritici blotch epidemics of winter wheat: build-up, overseasoning, and release of primary inoculum. Plant Pathology 60(2):166–177 (2011).

[27] Almogdad M, Lukošiūtė-Stasiukonienė A, Semaškienė R, Mačiulytė V. Sowing Date and Seed Rate Influence on Septoria Leaf Blotch Occurrence in Winter Wheat. Agriculture 14(7):988 (2024).

[28] AHDB. How drill date affects septoria disease resistance ratings in winter wheat (2025). URL https://ahdb.org.uk/news/how-drill-date-affects-septoria-disease-resistance-ratings-in-winter-wheat. [Online] (Accessed: 22 May 2025).

[29] Morais D, Sache I, Suffert F, Laval V. Is the onset of septoria tritici blotch epidemics related to the local pool of ascospores? Plant Pathology 65(2):250–260 (2016).

[30] Kristoffersen R, Jørgensen LN, Eriksen LB, Nielsen GC, Kiær LP. Control of Septoria tritici blotch by winter wheat cultivar mixtures: meta-analysis of 19 years of cultivar trials. Field Crops Research 249:107696 (2020).

[31] Borg J, Kiær LP, Lecarpentier C, Goldringer I, Gauffreteau A, et al. Unfolding the potential of wheat cultivar mixtures: A meta-analysis perspective and identification of knowledge gaps. Field Crops Research 221:298–313 (2018).

[32] Mikaberidze A, McDonald BA, Bonhoeffer S. Developing smarter host mixtures to control plant disease. Plant Pathology 64(4):996–1004 (2015).

[33] Clin P, Grognard F, Andrivon D, Mailleret L, Hamelin FM. Host mixtures for plant disease control: Benefits from pathogen selection and immune priming. Evolutionary Applications 15(6):967– 975 (2022).

[34] AHDB. Recommended Lists (2025). URL https://ahdb.org.uk/knowledge-library/recommended-lists-for-cereals-and-oilseeds-rl. [Online] (Accessed: 22 May 2025).

[35] Mann R, Kettlewell P, Jenkinson P. Effect of foliar-applied potassium chloride on septoria leaf blotch of winter wheat. Plant pathology 53(5):653–659 (2004).

[36] Perelló AE, Moreno MV, Mónaco C, Simón MR, Cordo C. Biological control of Septoria tritici blotch on wheat by Trichoderma spp. under field conditions in Argentina. BioControl 54:113–122 (2009).

[37] Kildea S, Ransbotyn V, Khan MR, Fagan B, Leonard G, et al. Bacillus megaterium shows potential for the biocontrol of Septoria tritici blotch of wheat. Biological control 47(1):37–45 (2008).

[38] DEFRA. Pest and Disease Survey (2025). URL https://www.pestanddiseasesurvey.co.uk/home. [Online] (Accessed: 22 May 2025).

[39] IPM Decisions Network. IPM Decisions (2025). URL https://www.ipmdecisions.net/. [Online] (Accessed: 22 May 2025).

[40] Kristoffersen R, Eriksen LB, Nielsen GC, Jørgensen JR, Jørgensen LN. Management of Septoria tritici blotch using cultivar mixtures. Plant Disease 106(5):1341–1349 (2022).

[41] Bekele E, Kassaye Z. Integrated management of Septoria blotches of wheat: Effect of sowing date, variety and fungicide. Pest Managment Journal of Ethiopia 7:11–18 (2003).

[42] Shaw M, Royle D. Airborne inoculum as a major source of Septoria tritici (Mycosphaerella graminicola) infections in winter wheat crops in the UK. Plant Pathology 38(1):35–43 (1989).

[43] Holmes S, Colhoun J. Straw-borne inoculum of Septoria nodorum and S. tritici in relation to incidence of disease on wheat plants. Plant Pathology 24(2):63–66 (1975).

[44] Dehbi I, Achemrk O, Ezzouggari R, El Jarroudi M, Mokrini F, et al. Beneficial Microorganisms as Bioprotectants against Foliar Diseases of Cereals: A Review. Plants 12(24):4162 (2023).

[45] Tang S, Xiao Y, Chen L, Cheke RA. Integrated pest management models and their dynamical behaviour. Bulletin of mathematical biology 67:115–135 (2005).

[46] Lane DE, Walker TJ, Grantham DG. IPM adoption and impacts in the United States. Journal of Integrated Pest Management 14(1):1 (2023).

[47] McQuaid CF, Gilligan CA, van den Bosch F. Considering behaviour to ensure the success of a disease control strategy. Royal Society Open Science 4(12):170721 (2017).

[48] Radici A, Martinetti D, Bevacqua D. Optimizing fungicide deployment in a connected crop landscape while balancing epidemic control and environmental sustainability. bioRxiv pages 2024–12 (2024).

[49] Tang S, Cheke RA. Models for integrated pest control and their biological implications. Mathematical Biosciences 215(1):115–125 (2008).

[50] Tang S, Tang G, Cheke RA. Optimum timing for integrated pest management: modelling rates of pesticide application and natural enemy releases. Journal of Theoretical Biology 264(2):623–638 (2010).

[51] KWS. Sowing wheat Information about seed rates, seed dates and soil preparation (2025). URL https://www.kws.com/gb/en/consulting/sowing/sowing-wheat/. [Online] (Accessed: 22 May 2025).

[52] DAERA Northern Ireland. Autumn drilling (2025). URL https://www.daera-ni.gov.uk/articles/autumn-drilling. [Online] (Accessed: 22 May 2025).

[53] Bellameche F, Pedrazzini C, Mauch-Mani B, Mascher F. Efficiency of biological and chemical inducers for controlling Septoria tritici leaf blotch (STB) on wheat (Triticum aestivum L.). European Journal of Plant Pathology 158(1):99–109 (2020).

[54] Le Mire G, Siah A, Brisset MN, Gaucher M, Deleu M, et al. Surfactin protects wheat against Zymoseptoria tritici and activates both salicylic acid-and jasmonic acid-dependent defense responses. Agriculture 8(1):11 (2018).

[55] Lynch K, Zannini E, Guo J, Axel C, Arendt E, et al. Control of Zymoseptoria tritici cause of septoria tritici blotch of wheat using antifungal Lactobacillus strains. Journal of Applied Microbiology 121(2):485–494 (2016).

[56] Steiro ÅL, Kvakkestad V, Breland TA, Vatn A. Integrated Pest Management adoption by grain farmers in Norway: A novel index method. Crop protection 135:105201 (2020).

[57] Creissen HE, Jones PJ, Tranter RB, Girling RD, Jess S, et al. Identifying the drivers and constraints to adoption of IPM among arable farmers in the UK and Ireland. Pest Management Science 77(9):4148–4158 (2021).

[58] Popp J, Peęo K, Nagy J. Pesticide productivity and food security. A review. Agronomy for sustainable development 33:243–255 (2013).

[59] Gisi U, Leadbeater A. The challenge of chemical control as part of integrated pest management. Journal of Plant Pathology pages S11–S15 (2010).

[60] Corkley I, Fraaije B, Hawkins N. Fungicide resistance management: Maximizing the effective life of plant protection products. Plant Pathology 71(1):150–169 (2022).

[61] Bottrell D, Schoenly K. Integrated pest management for resource-limited farmers: challenges for achieving ecological, social and economic sustainability. The Journal of Agricultural Science 156(3):408–426 (2018).

[62] Corkley I, Helps J, van den Bosch F, Paveley ND, Milne AE, et al. Delaying Infection Through Phytosanitary Soybean-Free Periods Contributes to Fungicide Resistance Management in Phakopsora pachyrhizi: A Modelling Analysis. Plant Pathology (2025).

[63] Semenov MA. Impacts of climate change on wheat in England and Wales. Journal of the Royal Society Interface 6(33):343–350 (2009).

[64] Bajwa AA, Farooq M, Al-Sadi AM, Nawaz A, Jabran K, et al. Impact of climate change on biology and management of wheat pests. Crop Protection 137:105304 (2020).

[65] Gvozdenac S, Dedić B, Mikić S, Ovuka J, Miladinović D. Impact of climate change on integrated pest management strategies. Climate Change and Agriculture: Perspectives, Sustainability and Resilience pages 311–372 (2022).

[66] Zayan SA. Impact of climate change on plant diseases and IPM strategies. In Plant diseases-current threats and management trends. IntechOpen (2019).

