## Supporting Information for "Modelling the effectiveness of Integrated Pest Management strategies for the control of Septoria tritici blotch"

#### Table of Contents

**S1 Base model** 2

**S2 Variation in optimal sowing date** 3

**S3 Additional figures** 4

### S1 Base model

Our base model for the infectious disease dynamics of Septoria tritici blotch (STB) (in the absence of any controls) is adapted from the system of ordinary differential equations (ODEs) in Hobbelen *et al.* [1], with up-to-date parameter values taken from Corkley *et al.* [2]. A schematic representation of the system is shown in Fig. 1.

In the absence of any controls, the following system of ODEs describes infection with STB:

$$
\begin{aligned}
\frac{dS}{dt} &= g(A,t) - \sigma(t)S - \frac{\beta S(I+P)}{A} \\
\frac{dE}{dt} &= \frac{\beta S(I+P)}{A} - \sigma(t)E - \gamma E \\
\frac{dI}{dt} &= \gamma E - \mu I \\
\frac{dR}{dt} &= \sigma(t)(S+E) \\
\frac{dD}{dt} &= \mu I \\
\frac{dP}{dt} &= -\nu P.
\end{aligned} \tag{7}
$$

Here $\sigma(t)$ represents the rate of senescence of healthy tissue [1, 2]

$$
\sigma(t) = \begin{cases} 0 & \text{if } t < T_{61} \\ 0.0028\frac{t-T_{61}}{T_{87}-T_{61}} + 0.704e^{-0.314*(T_{87}-t)} & \text{if } T_{61} \leq t, \end{cases} \tag{8}
$$

and $g(A,t)$ represents the growth of healthy tissue [1] and was defined by

$$
g(A,t) = \begin{cases} 0 & \text{if } A = 0 \\ G(A_{\max} - A) & \text{otherwise.} \end{cases} \tag{9}
$$

Note that we have included the condition that $g(A,t) = 0$ when $A = 0$ while Hobbelen *et al.* [1] do not. We specified this additional healthy tissue growth condition so that late sowed fields do not experience growth until the initial condition is explicitly set to be non-zero.

The initial conditions, at the beginning of a year's growing season, were $(S,E,I,R,D,P) = (0.05,0,0,0,0,\Psi)$ [1–3].

For a summary of model parameters and their values, see Table 1.

#### S2 Variation in optimal sowing date

Note that when the full low-, medium- and high-intensity IPM regimes are defined, most values are taken directly from the low, medium and high values of each individual control measures. However, the sowing date which optimises yield differs when used in combination with other IPM measures, compared to when it is used as the only control measure. This occurs because of the trade-off in yield which comes from delayed sowing; later sowing results in the evasion of disease, but a lower density of (healthy) tissue during the grain forming period. But as the presence of other control methods affects disease prevalence, it also affects which sowing date will optimise the trade-off between disease evasion and grain development. Fig. S1 shows how the optimal sowing date varies depending on the other active controls.

Based on the outputs from Fig. S1, we maintained the 7 day delay as a medium-effectiveness value alongside $q = 0.154$ for the medium-intensity regime. But we changed the sowing delay in the high-intensity IPM regime, updating it from the 14 day delay which optimised yield when this intervention is in isolation, to a 3 day delay which optimised the yield when delayed sowing is implemented alongside other interventions.

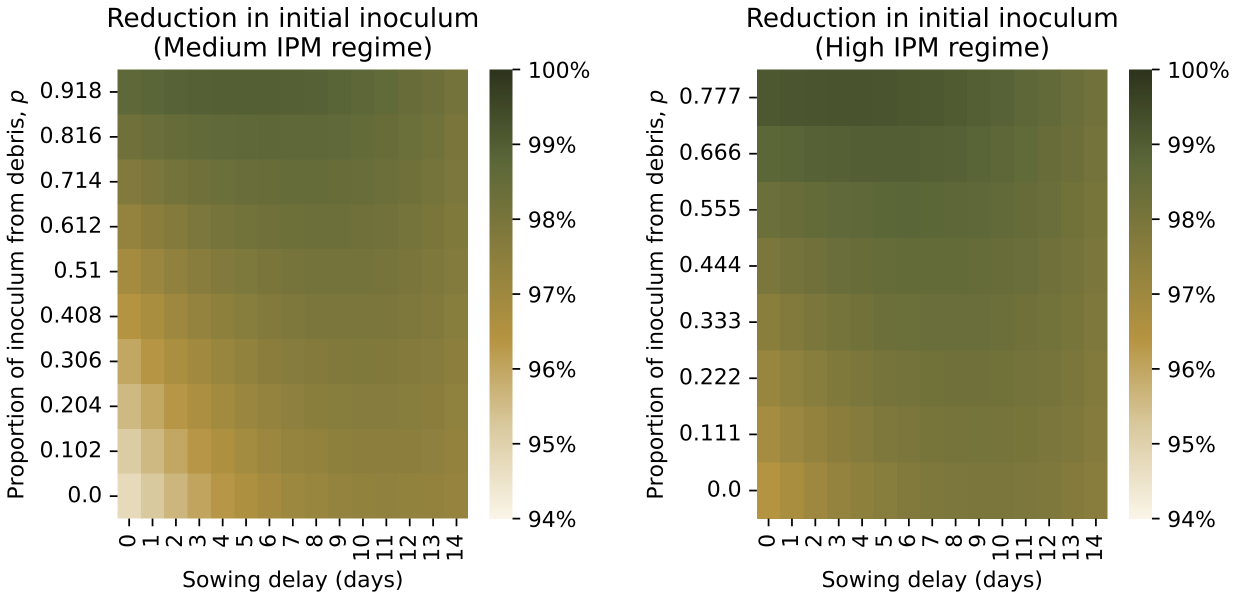


**Fig. S1. Percentage of initial inoculum ($\Psi$) which remains at the start of the season.** Heatmaps showing the proportion of initial inoculum which remains at the start of the season, as $p$ and the number of days sown late varies. Left panel shows the outcomes in the medium-regime scenario ($r_\beta = 0.985$, biocontrol applied at $T_{31}$, $q = 0.154$). Right panel shows the outcomes in the high-regime scenario ($r_\beta = 0.972$, biocontrol applied at $T_{31}$ and $T_{31,39}$, $q = 0$). The sowing delay which results in the highest disease reduction varies, both across the values of $p$ in each scenario, and between scenarios.

#### S3 Additional figures

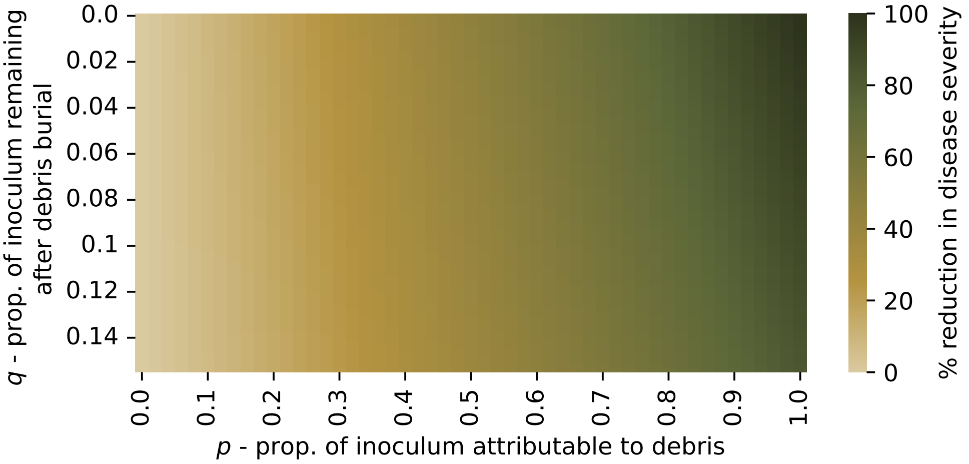


**Fig. S2. Reduction in disease severity resulting from pairs of $p$ and $q$ values.** The percentage reduction in disease severity for all pairs of $p$ between 0 and 1 and $q$ between 0 and 0.154 are shown. Darker shading corresponds to a greater reduction in disease severity. For fixed $q$, we see that a different range of $p$ is required to fit for a reduction of between 0-75% disease severity.

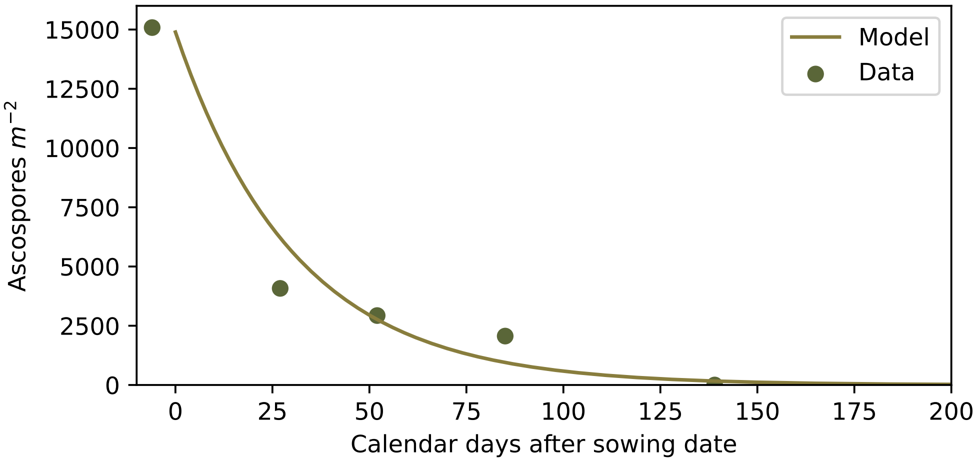


**Fig. S3. Model fit to ascospore data from Morais *et al.* [4].** Model density of ascospore prevalence (approximate number per $m^2$), dependant on the number of calendar days since the sowing date. Model density was fitted to data from plots with debris [4]. Data is shown as dark green circles, and the fit function is shown in the lighter green line.

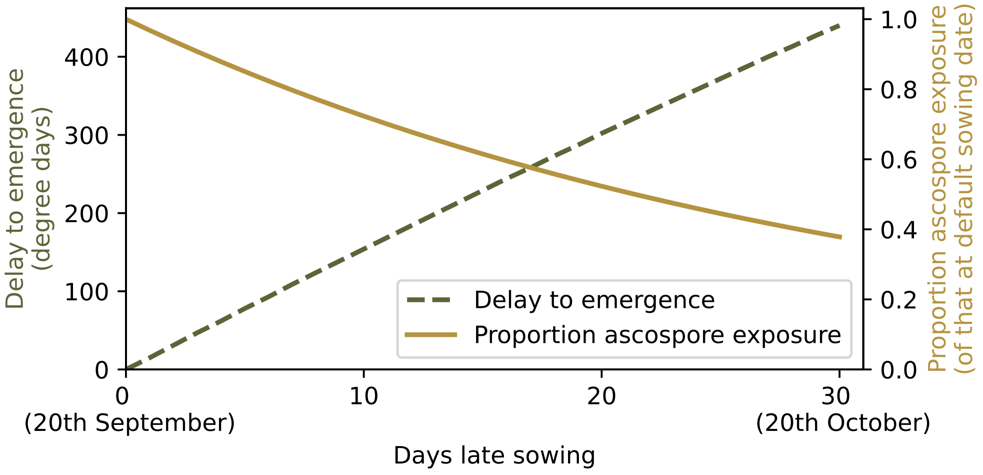


**Fig. S4. Ascopore exposure and emergence dependencies on sowing delays.** Left y-axis shows the level of ascospore exposure for a range of sowing delays (dashed line). Right y-axis shows the degree day delay to emergence for a range of sowing delays (solid line).

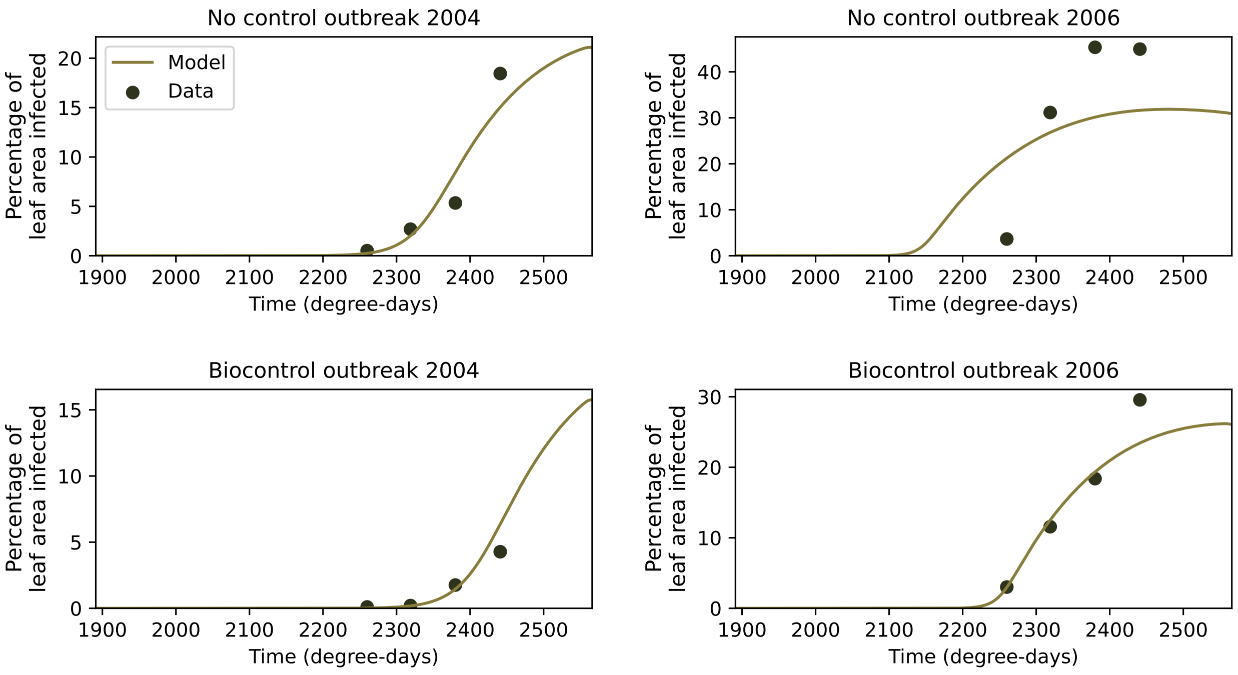


**Fig. S5. Model fit to 2004 and 2006 experimental infection data from Kildea *et al.* [5].** Data points are shown as filled circles and model fits are shown as solid lines. We first performed fitting on the 'no control' data to determine the outbreak severity in 2004 and 2006 (top row). Using each year's respective severity parameter, to find the parameters for biocontrol in our model we then performed fitting to the data from the plants with biocontrol applied (bottom row).

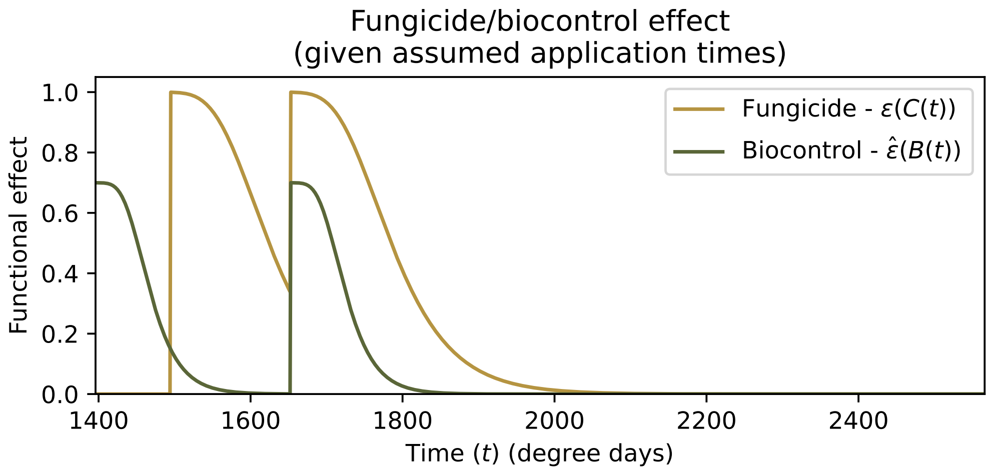


**Fig. S6. Comparison between the functional outputs of the concentration of fungicide (Eq. (2)) and the concentration of biocontrol treatment (Eq. (4)).** For the displayed results for the biocontrol treatment we assumed there is an application at $T_{31}$ and at $T_{39}$.

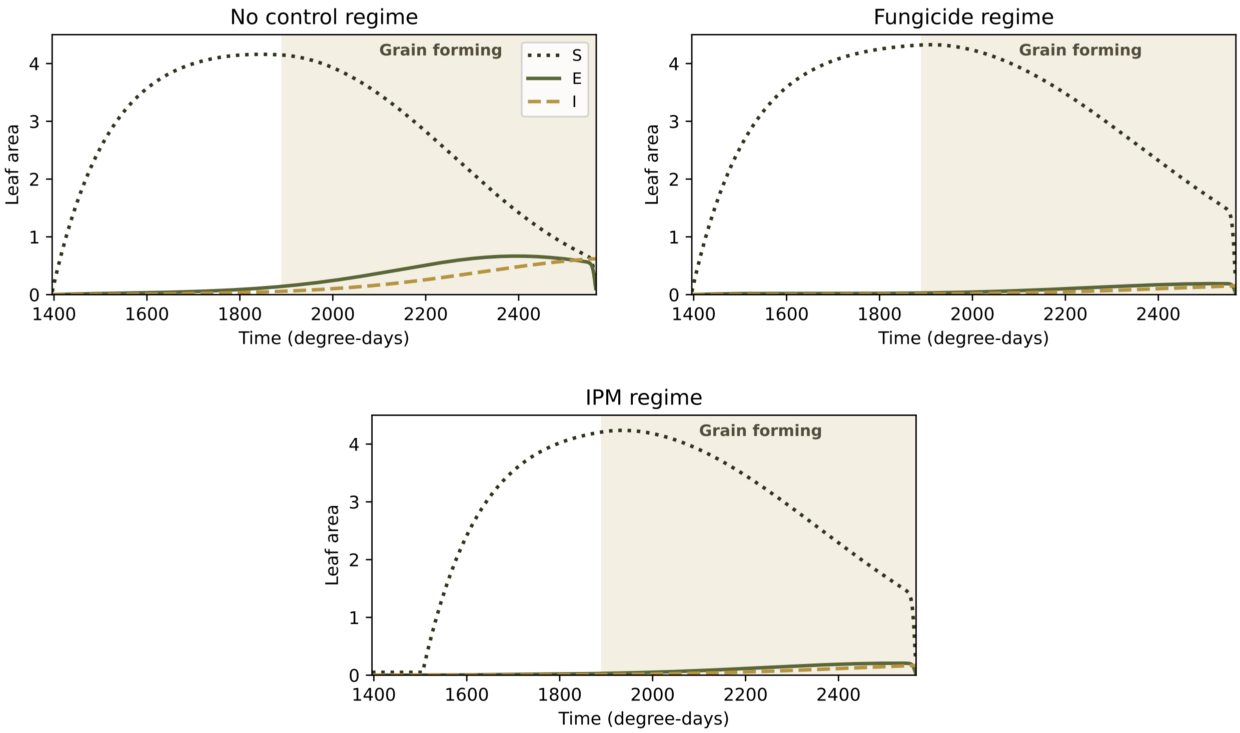


**Fig. S7. Outbreak in farms under three different regimes: No control, fungicide, and IPM.** Temporal progression of outbreak is shown in each farm type, by the total leaf area in compartments $S$ (black dotted lines), $E$ (solid green lines) and $I$ (yellow dashed lines).

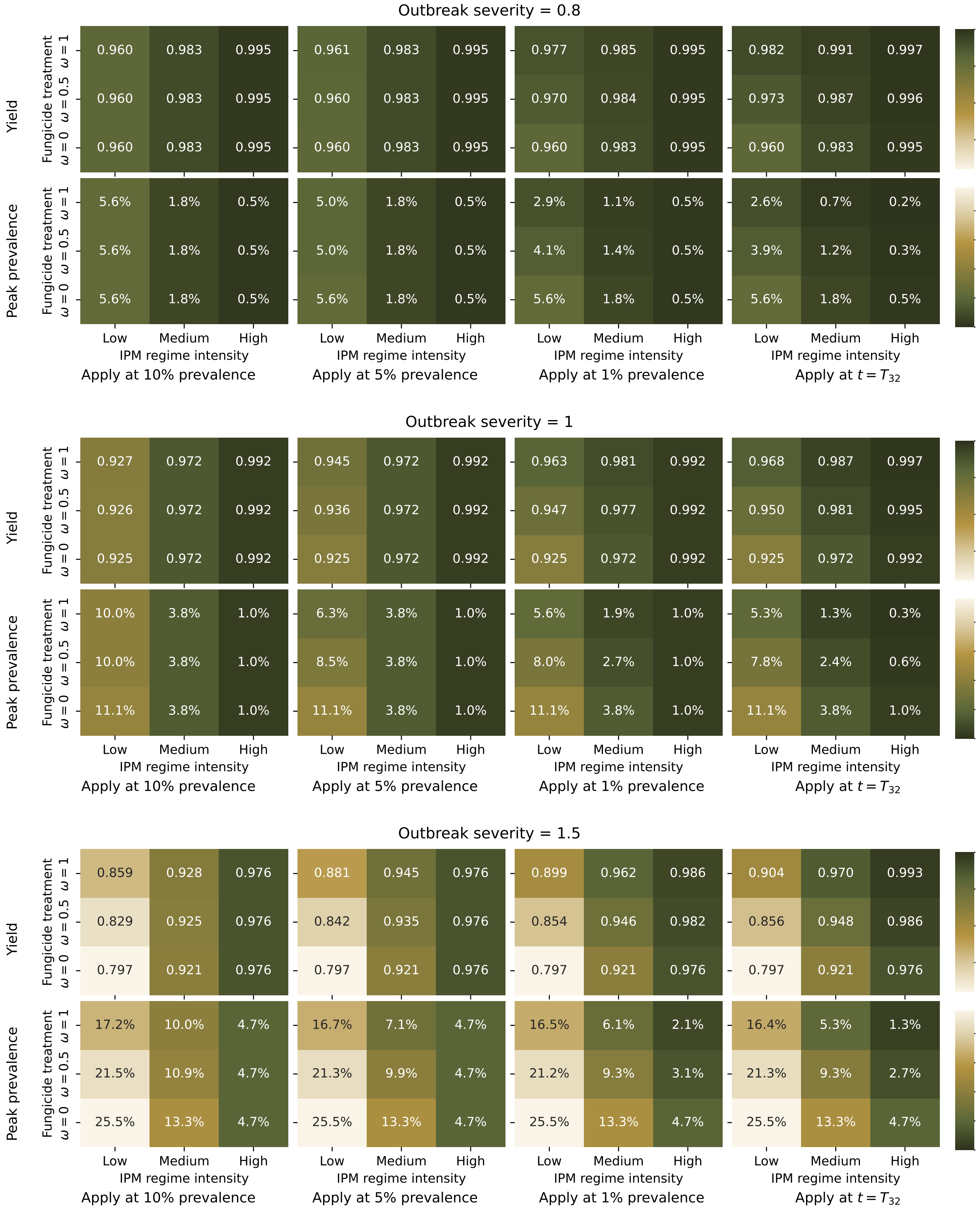


**Fig. S8. Yield and peak infection prevalence outcomes under varying environmental severity and considering the use of reactive fungicide treatments.** The resulting yield and peak prevalence in three outbreak severities; low-severity (0.8) (top panel), medium/standard-severity (1) (center panel) and high-severity (1.5) (bottom panel). Each panel contains the results under four reactive fungicide treatment regimes; application at $t = T_{32}$, at 1%, 5% and 10% prevalence. Each fungicide treatment regime considers the application of a full ($\omega = 1$) and reduced ($\omega = 0.5$) fungicide dose, as well as the case with no fungicide applied ($\omega = 0$) for comparison. The results are found in all three of our considered IPM regimes; low-, medium- and high-intensity.

#### Supplementary references


[1] Hobbelen P, Paveley N, Van den Bosch F. Delaying selection for fungicide insensitivity by mixing fungicides at a low and high risk of resistance development: A modeling analysis. *Phytopathology* **101**(10):1224–1233 (2011).

[2] Corkley I, Mikaberidze A, Paveley N, van den Bosch F, Shaw MW, *et al.* Dose Splitting Increases Selection for Both Target-Site and Non-Target-Site Fungicide Resistance—A Modelling Analysis. *Plant Pathology* (2025).

[3] Elderfield JA, Lopez-Ruiz FJ, van den Bosch F, Cunniffe NJ. Using epidemiological principles to explain fungicide resistance management tactics: Why do mixtures outperform alternations? *Phytopathology* **108**(7):803–817 (2018).

[4] Morais D, Sache I, Suffert F, Laval V. Is the onset of septoria tritici blotch epidemics related to the local pool of ascospores? *Plant Pathology* **65**(2):250–260 (2016).

[5] Kildea S, Ransbotyn V, Khan MR, Fagan B, Leonard G, *et al.* Bacillus megaterium shows potential for the biocontrol of Septoria tritici blotch of wheat. *Biological control* **47**(1):37–45 (2008).
